# Inferring metabolic rewiring in embryonic neural development using single cell data

**DOI:** 10.1101/2020.09.03.282442

**Authors:** Shashank Jatav, Saksham Malhotra, Freda D Miller, Abhishek Jha, Sidhartha Goyal

## Abstract

Metabolism is intricately linked with cell fate changes. Much of this understanding comes from detailed metabolomics studies averaged across a population of cells which may be composed of multiple cell types. Currently, there are no quantitative techniques sensitive enough to assess metabolomics broadly at the single cell level. Here we present *scMetNet*, a technique that interrogates metabolic rewiring at the single cell resolution and we apply it to murine embryonic development. Our method first confirms the key metabolic pathways, categorized into bioenergetic, epigenetic and biosynthetic, that change as embryonic neural stem cells differentiate and age. It then goes beyond to identify specific sub-networks, such as the cholesterol and mevalonate biosynthesis pathway, that drive the global metabolic changes during neural cortical development. Having such contextual information about metabolic rewiring provides putative mechanisms driving stem cell differentiation and identifies potential targets for regulating neural stem cell and neuronal biology.

## Introduction

The rapid development of computational methods for single cell gene expression has led to new insights into developmental biology. These insights range from confirmation of known knowledge accumulated over decades from “one” experiment (Farrell et al., 2018; Karaiskos et al., 2017), to discovering new cell types and their lineages (Kester & van Oudenaarden, 2018) and addressing tumor heterogeneity (Levitin, Yuan, & Sims, 2018). However, many challenges remain (Lähnemann et al., 2020) creating as many opportunities (Dasgupta, Bader, & Goyal, 2018).

One of the fundamental challenges is that while the cell types in developing tissues can be discerned the identity of key regulators behind the emergence of these cell types are hard to pinpoint. This is due to the low expression levels and often transient nature of transcription factors (Martin & Sung, 2018) that are thought to drive cell transitions. Another limitation of understanding key regulators from single cell analysis is not knowing the landscape of interactions between proteins, genes, and small molecules. This is particularly challenging as these interactions change with biological context.

In addition to master gene regulators such as transcription factors, metabolic rewiring is known to accompany and regulate changes in cell state. For example, metabolism is being targeted in multiple disease states (Afonso, Santos, Longatto-Filho, & Baltazar, 2020; Huang & Perl, 2018; Luengo, Gui, & Vander Heiden, 2017). As a second example, metabolic rewiring accompanies early development (Intlekofer & Finley, 2019; Shyh-Chang & Ng, 2017) and external metabolites have been used to alter developmental output in organoids (Schell et al., 2017), suggesting a potential for effective *in vivo* metabolic interventions in stem cell biology.

Metabolism has been studied extensively within stem cells, with a focus on *in vitro* study of mouse embryonic stem cells (mESCs) and induced pluripotent stem cells (iPSCs) as biological models (Shyh-Chang & Ng, 2017). These studies have defined various metabolic pathways that govern stem cell fate decisions, proliferation and differentiation. Can single cell sequencing shed light on metabolic regulators of *in vivo* development? Prima facie, in the case of metabolism, two of the main challenges associated with understanding regulators of stem cell fate dynamics, low expression levels of transcription factors and lack of interaction map between the signaling proteins, may not be limiting. Genes that encode metabolic enzymes are often expressed at high levels, and a detailed understanding of metabolic pathways provides the required interaction landscape. Recently, metabolic changes in tumors have been addressed using single cell expression data (Xiao, Dai, & Locasale, 2019), thereby demonstrating the potential importance of such an approach at the therapeutic level.

Here we present *scMetNet*, a method that uses single cell gene expression data to create a map for metabolic rewiring between different cell types at a particular stage and for a particular cell type across developmental time. We have applied *scMetNet* to understand embryonic neural development, with a particular focus on the murine embryonic cortex from E13.5 to E17.5, a developing tissue that is comprised of radial glial precursors (RPs) that generate neurons either directly or via intermediate progenitors (IP), a neurogenic transit-amplifying cell (Yuzwa et al., 2017). Using this approach, we define unique metabolic states that are aligned significantly with RPs, IPs and newborn neurons. Given the significant concordance between cell types and their metabolism, we then inferred a metabolic network based on the known landscape of metabolic interactions from the Kegg database (Kanehisa & Goto, 2000). This connected network of genes and metabolites defined the metabolic re-wiring between RPs and neurons within each time point. As neurons increase in number to form the cortical networks that serve higher cognitive functions, RPs alter their metabolic character as they switch from generating neurons to transitioning to postnatal dormant forebrain neural stem cells. We document a similar metabolic signature using single cell gene expression from a second region of the brain where embryonic neural precursors persist to become dormant postnatal neural stem cells, the embryonic hippocampal dentate gyrus. Overall, scMetNet identified the precise metabolic rewiring that occurs as embryonic neural stem cells generate neurons, and provides potential molecular targets for regulating neural development.

## Results

Here, we utilized scMetNet and analyzed single cell gene expression data from the murine embryonic cortex from E13.5 to E17.5, when there are three distinct cell types: radial glial precursors (RPs), a transit-amplifying intermediate progenitor cell called an intermediate progenitor (IP) and their newborn neuronal progeny (Figure 1A).

**Figure 1:**
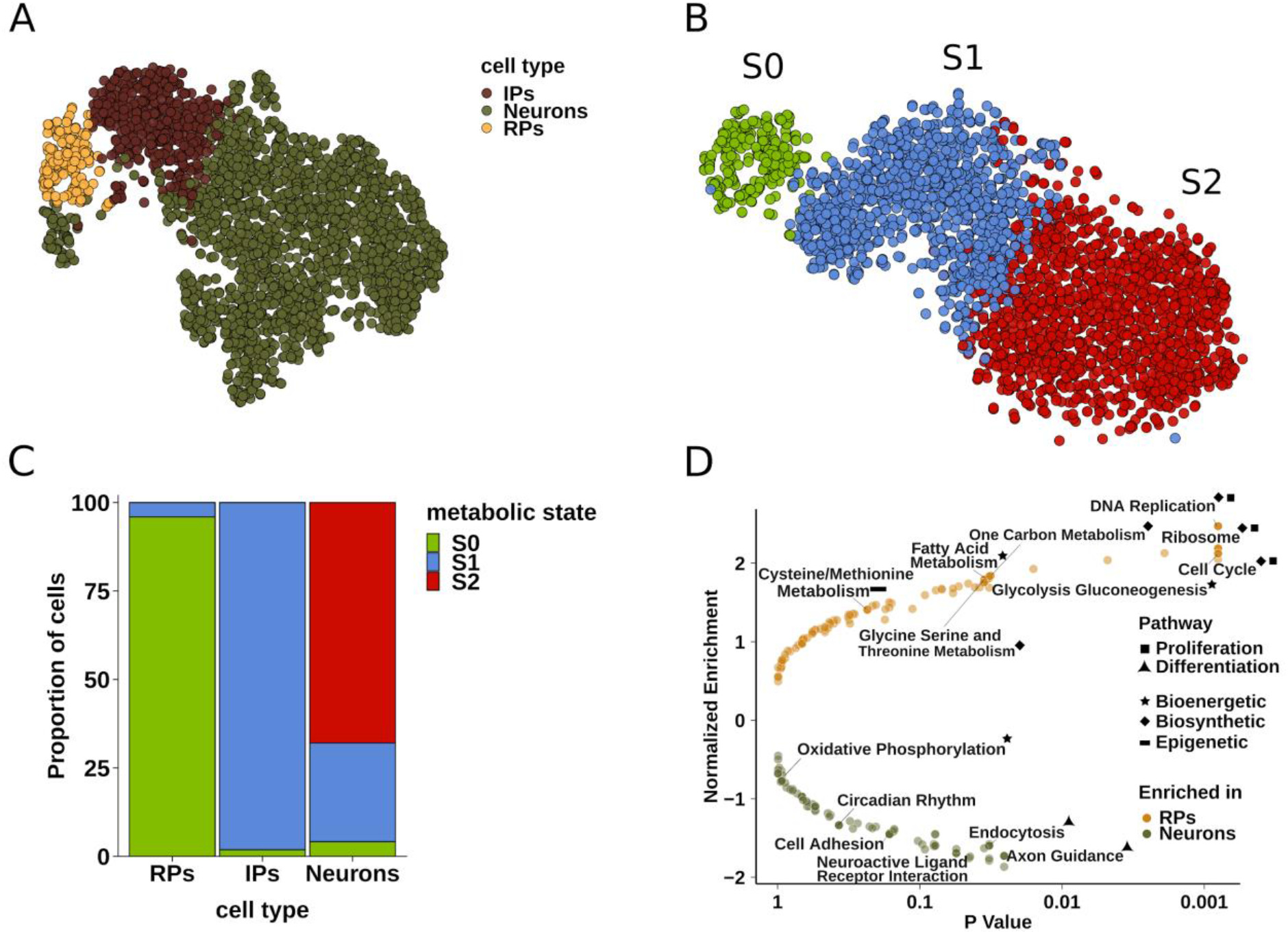
Cell types align with metabolic states. (A) t-distributed stochastic neighbour embedding (t-SNE) visualizations of cell types at E15.5. The clustering was done using highly variable genes, cell type-specific markers for Neurons, IPs and RPs were used to annotate cells. (B) t-SNE visualizations of metabolic states at E15.5. The clustering was done using only the metabolic genes. (C) The overlap of cell types and metabolic states at E15.5; here RPs are predominantly in metabolic state S_0_ and all IPs are present in metabolic state S_1_. (D) Gene Set Enrichment Analysis (GSEA) between RPs and Neurons shows the enriched pathways in the two cell types.

### Metabolic states align with developmental lineages in embryonic neural development

To understand the metabolic changes that hallmark the neural stem cell to neuron transition during embryonic cortical development, we asked whether there are distinct metabolic states that accompany emerging cell types. To address this question, cells were clustered using only their metabolic gene expression profiles (Fig. 1B and Methods Section for details). These clusters defined the “metabolic states” of the cells in the developing cortex. Notably, by delineating the induced transcriptional metabolic pathway changes, three distinct metabolic states emerged that aligned well with the three embryonic cortical cell types (Fig. 1C). Can this alignment result from chance? To test this, we computed the overlap of cell types with clusters derived from random sets of non-metabolic genes with each set having the same number of genes as the metabolic genes. We found that there are no random sets that are better aligned with RPs than metabolic genes, while there may be some random sets that can show higher alignment with neurons and IPs. Overall, metabolic genes show significant overlaps (pvalue <10^-3^) across all the three cell types (Supplementary Fig. S1).

The metabolite specific alignment details a compelling picture of the temporal metabolic changes that accompany the transition between neural precursors and neurons. To determine the effect of number of metabolic states, we computed overlap of cell types with increasing numbers of metabolic clusters. We found that the largest cluster of cells annotated as neurons gets further segmented into different metabolic states (see Supplementary Fig. S2), consistent with multiple cell clusters observed earlier using all the variable genes (Yuzwa et al. 2017). Overall, these metabolic states were stably represented across the three developmental time points considered (see Supplementary Fig. S1). In particular at E15.5, the majority of cells identified as RPs map to metabolic state S0, most of the IPs are in metabolic state S1, and neurons are in S1 and S2. The temporal change in the metabolic state of IPs is consistent with their transient nature; they first were almost equally split between metabolic states S0 and S1 at E13.5, then at E15.5 they were almost fully in state S1 and then they were no longer present at E17.5. This is notable, as by E17.5 RPs are known to be generating very few neurons, hence may not be generating IPs (shown in Supplementary Fig. S1).

### The metabolic changes between RPs and neurons reflect classic hallmarks of cellular differentiation

Given that the three cell types are largely in different metabolic states, we applied Gene Set Enrichment Analysis (GSEA) to ask what pathways separate the undifferentiated RPs from the differentiated neurons. GSEA leverages annotated pathway databases to identify “enriched” pathways by comparing the average gene expression between two cell types (Fig. 1D). Consistent with the fact that RPs proliferate but neurons are post-mitotic, the most enriched gene sets in RPs are those involving DNA replication and the cell cycle. For neurons, the most significantly enriched are axon guidance, and neuronal migration-related genes.

When focusing on the metabolic subsets, GSEA confirmed important pathways previously seen during cellular differentiation in other systems. Many of these pathways fall into three functional categories often used to describe metabolic changes during tissue development (Shyh-Chang & Ng, 2017). First are bioenergetic pathways necessary to support rapid cell division in developing tissues. As such fatty acid oxidation and glycolysis were upregulated in RPs suggesting their role in meeting the energy requirements of these rapidly dividing cells. Second are epigenetic pathways involved in the large-scale epigenetic changes that drive cellular differentiation. Here, methionine metabolism was upregulated presumably to supply the methylation demands associated with modifying the epigenome of RPs differentiating into neurons (Shiraki et al., 2014). Third are biosynthetic pathways necessary for the rapid growth of biomass necessary to sculpt tissues. Here, we found glycine-serine and one carbon metabolism were the biosynthetic pathways driving synthesis of macromolecules, such as glycine and serine, essential for increasing neuron numbers during brain development. A restricted GSEA analysis only for metabolic pathways applied on the cells in the different metabolic states was consistent with the RP-vs-neurons GSEA analysis (Supplementary figure S3).

### scMetNet reveals inter-connected metabolic modules

While the GSEA analysis identifies average changes that occur in different cell types during differentiation, it does not reveal specific genes that define these significant metabolic pathways, the specific reactions they catalyze, and/or the metabolites that are important. Having such a finer scale understanding is essential to altering the fate of cells and developing new potential therapeutics. We addressed these issues by adapting a network algorithm, CombiT (Jha et al., 2015), designed originally to utilize microarray and metabolite measurement data, into scMetNet pipeline to evaluate for metabolic rewiring that differentiates RPs from neurons. Specifically, CombiT algorithm allows scMetNet to identify the most connected network consisting of enzyme-metabolite reactions using the Kegg database that define the difference between the RPs and neurons. In addition to metabolic gene networks, scMetNet leverages the curated metabolic knowledge to infer the critical metabolites that form the metabolic backbone of the re-wired network. (Refer to methods)

scMetNet analysis of the E15.5 cortical cells (shown in Fig. 2) highlights differences in key genes and inferred metabolites that distinguish metabolism between RPs and neurons. As shown, the full metabolic network can be easily split into interconnected modules that align with the three functional categories of GSEA derived significant pathways, bioenergetic, epigenetic and biosynthetic.

**Figure 2:**
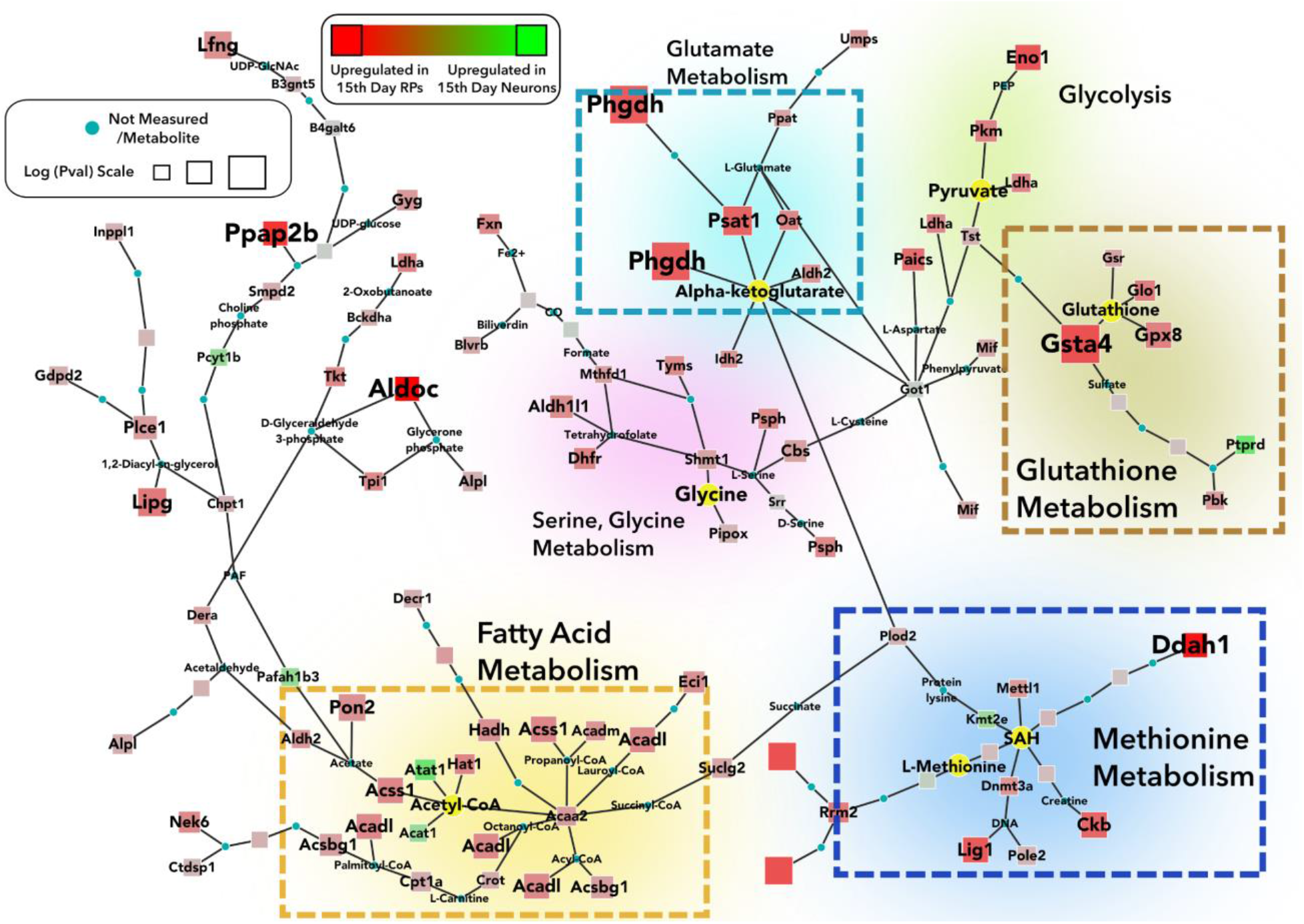
scMetNet network between RPs vs Neurons at E15.5. The circular nodes represent metabolites that connect the metabolic enzymes represented by square nodes; the size of square nodes around the enzymes denotes significance. Significant metabolic modules (in dashed boxes), and important metabolites (in yellow) are highlighted.

With regard to bioenergetics, upregulation of both fatty acid and glycolysis modules in RPs support metabolism-driven oxidative-phosphorylation (OXPHOS) to fulfill the high-energy requirements of rapidly dividing RPs. Within the fatty acid module, surprisingly, both biosynthetic and catabolic genes were upregulated. In particular, upregulation of genes like *Acsbg1*, which is involved in the synthesis of long chain fatty acids, suggest a biosynthetic role of fatty acid metabolism while upregulation of *Cpt1a*, which transports fatty acids to mitochondria, suggests a bioenergetic role utilizing glycolysis (Stephens, Constantin-teodosiu, & Greenhaff, 2007). More directed approaches will be required to distinguish futile metabolic cycling reactions versus activation of different subnetworks for some other process.

Although the OXPHOS module is not significantly different between RPs and neurons (Fig. 1D), its importance for RPs is suggested from the observed upregulation of *Psat1* and *Phgdh* which influence intracellular levels of alpha ketoglutarate (a-KG) (Hwang et al., 2016; Reid et al., 2018), an important component in the TCA cycle. The importance of the TCA cycle is also supported by upregulation of *Idh2*. Thus, RPs seem to utilize a mix of glycolysis and OXPHOS for their energy requirements. It is worth noting that while the GSEA did not identify OXPHOS as an enriched pathway, scMetNet pipeline revealed parts of OXPHOS pathway that are connected to other modules such as glycolysis, and hence might be important to the overall metabolic rewiring at this stage of development.

The second functional change involves epigenetic pathways, which center around the two key metabolites, alpha-ketoglutarate (a-KG) and methionine. In this regard, a-KG, a key cofactor for the TET family of DNA hydroxylases and Jumonji C-domain-containing histone demethylases, is linked with enzymes *Psat1* and *Phgdh*. Notably, these genes are thought to determine the timing of differentiation of embryonic stem cells by regulating a-KG levels (Hwang et al., 2016), and these findings suggest they may play a similar role in RPs. Methionine metabolism and methylation reactions are also apparently upregulated in RPs, as indicated by genes such as *Dnmt3a* and *Mettl1*, which likely play a role in maintaining SAM levels (Shiraki et al., 2014). The observed upregulation of specific genes controlling levels of a-KG and methionine metabolism suggest potential new targets for altering the neural stem cell to neuron transition at the epigenetic level.

With regard to the third functional category, biosynthesis, enzymes driving glycine/serine metabolism and one carbon metabolism provide rapid synthesis of macromolecules as they are required in fast dividing RPs. Specifically, the scMetNet network (Fig. 2) identifies upregulation of genes essential for anabolic reactions that are part of the Phosphate Pentose Pathway (PPP) such as *Tpi1* and *Tkt*, as well as for synthesis of a variety of macromolecules from nucleotides (*Shmt1, Paics*) to fatty acids (*Acsbg1*) (Stincone et al., 2015). It also identifies links between different modules, where one carbon metabolism influences methionine metabolism, and glycine and serine metabolism feed into one carbon metabolism by acting as a source for folate intermediates (Amelio, Cutruzzolá, Antonov, Agostini, & Melino, 2014). Specific enzymes, such as *Tpi1* and *Tkt* in PPP, *Mthfd1, Dhfr*, and *Tyms* in one carbon metabolism, and *Psph, Shmt1* and *Paics* in glycine-serine metabolism provide particular targets for altering RP neuronal output at this stage.

### Cortical radial precursors shift their metabolic state as they transition from rapidly-proliferating to slowly-proliferating/quiescent during embryogenesis

While cortical RPs largely maintain their transcriptional identity over the developmental period from E13.5 to E17.5 (Yuzwa et al., 2017), they undergo a poorly-understood transition from rapidly-proliferating to slowly-proliferating as they switch to making glial cells and to populating the adult neural stem cell pool. We asked if this transition also involved metabolic changes, initially using differential GSEA. This analysis identified methionine, glycolysis and fatty acid metabolism as the most significantly changing pathways in RPs from E13.5 to E17.5 (Table S2 and S3). Methionine metabolism and glycolysis are downregulated at later developmental times, while fatty acid metabolism is significantly upregulated as shown by changes in gene enrichment scores for these pathways (Fig. 3A).

**Figure 3:**
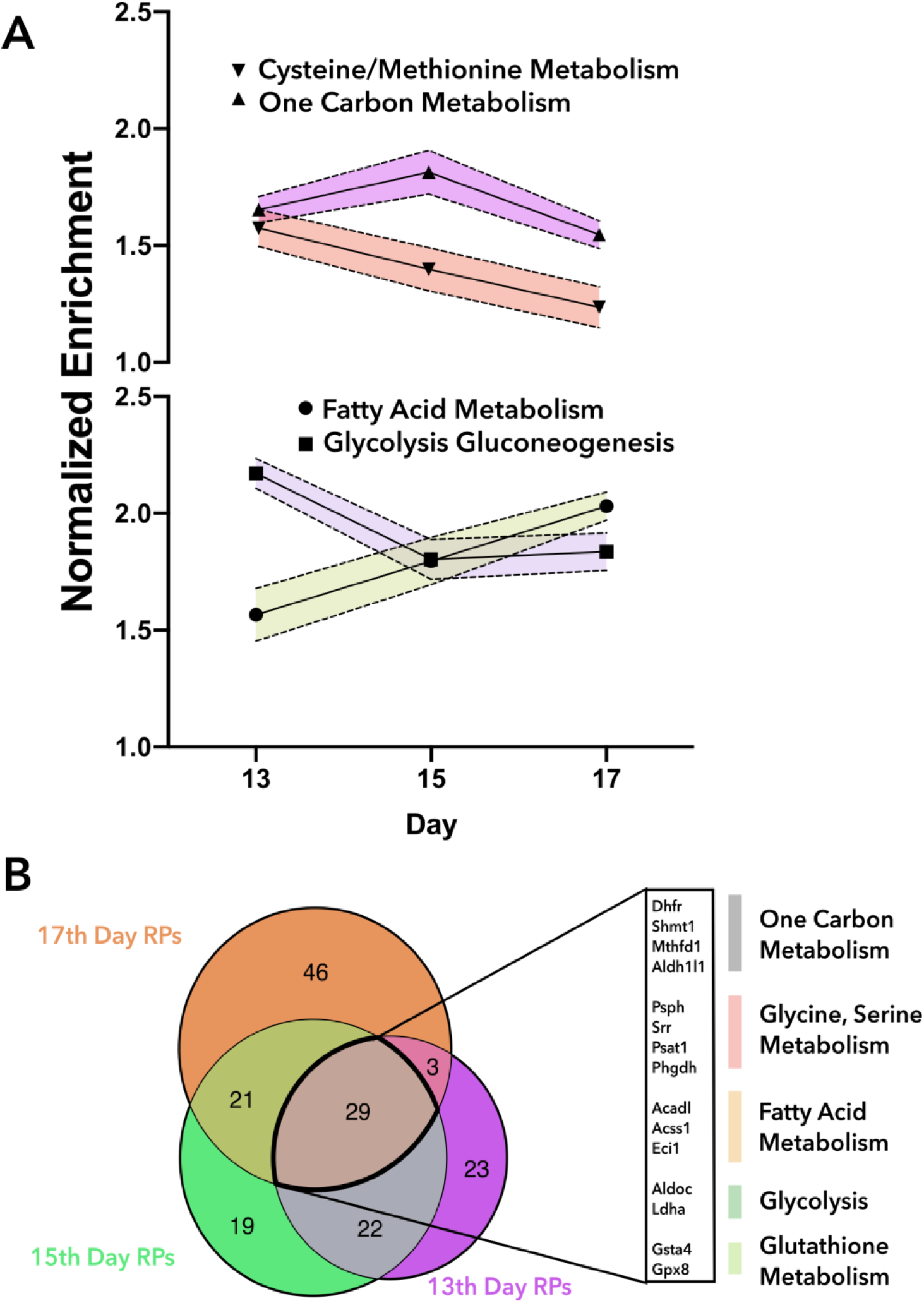
Metabolic ageing in radial precursors (RPs). (A) Global pathway changes between RPs and neurons using Gene Expression Enrichment Analysis (GSEA) at E13.5, E15.5 and E17.5 stages of embryonic development. (B) The core metabolic genes across E13.5, E15.5 and E17.5 and their associated pathways are identified.

These temporal trends are reflected in gene expression of key markers for these pathways that are assembled by the network analysis at the three time points (Fig. 3B shows the genes that are assembled by scMetNet and networks for E13.5 and E17.5 are shown in see Supplementary Fig. S4 and S5). However, while there are changes from E13.5, there are twenty-nine common metabolic genes that remain significant across the different time points and thus form the core metabolism that define RPs (see highlighted genes in Fig. 3B). We also observe upregulation of fatty acid and lipid synthesis at E15.5 and E17.5 (see Supp Fig S6). The core metabolism genes in general correspond to glycine-serine and one carbon metabolism highlighting the importance of these pathways in neuronal development (Supp Fig S6 and Fig 3B).

Notably, over this timeframe, based on the increase in fatty acid metabolism, which we had suspected is feeding into TCA cycle, and decrease in glycolysis in RPs we hypothesize an apparent preference for OXPHOS over glycolysis, which may suggest an increasing role for ROS signaling in RP differentiation, as previously suggested (Madhavan, Ourednik, & Ourednik, 2006).

To examine these similarities and differences more closely, we performed the scMetNet analysis of RPs from E13.5 and E17.5 (see Fig. 4 for the metabolic network). Consistent with the global analysis, the metabolic rewiring network showed an upregulation of fatty acid metabolism that involved genes involved in both biosynthesis (*Acsbg1*) and beta oxidation (*Acadl, Acaa2, Hadh*). Notably, *Cpt1a*, which was identified as likely being important for fatty acid metabolism driving OXPHOS, is not picked up by the scMetNet pipeline, suggesting that there is no significant shift in metabolic rewiring around *Cpt1a*. Instead, the upregulation of fatty acid synthesis between E13.5 and E17.5 involves a significant upregulation of cholesterol and mevalonate biosynthesis pathways, with the mevalonate biosynthesis pathway feeding directly into cholesterol synthesis through squalene.

**Figure 4:**
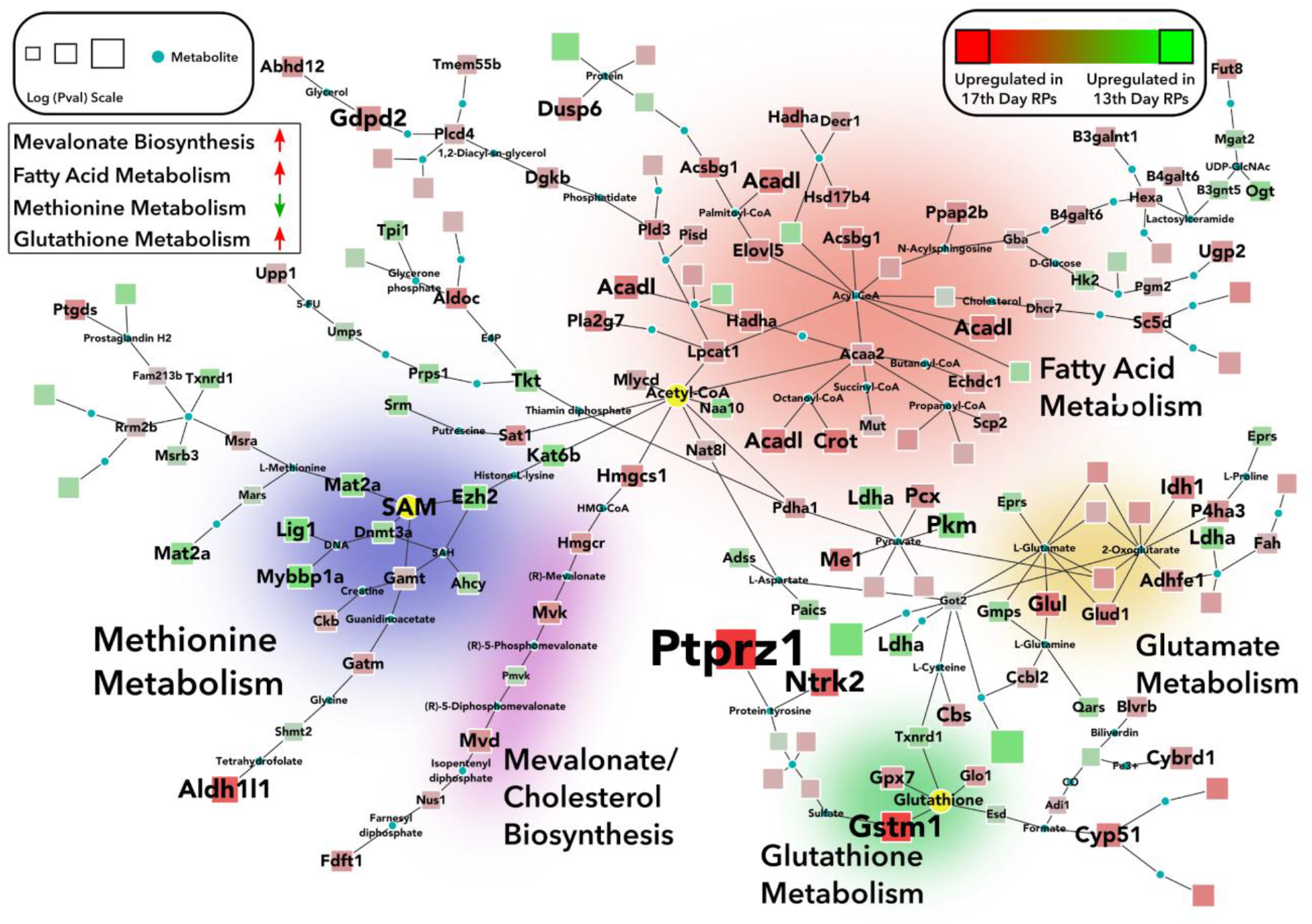
scMetNet network between E17.5 and E13.5 RPs show how metabolism gets re-wired within RPs along developmental time; the circular nodes represent metabolites that connect the metabolic enzymes represented by square nodes; the size of square nodes around the enzymes denotes significance. The mevalonate biosynthesis, fatty acid and glutathione pathways are upregulated while the methionine metabolism is downregulated at E17.5.

Notably, consistent with our hypothesis about the increasing role of ROS based on the global changes, scMetNet network (Fig. 4) shows glutathione S transferases and peroxidases (Gstm1, Gpx7) as being upregulated in later stage RPs. Taking a clue from the literature, this suggests that perturbing RPs ability to use peroxidases may increase neural differentiation and help alleviate pathologies related to delayed neural development (Savaskan, Borchert, Bräuer, & Kuhn, 2007). Here again CombiT revealed aspects of metabolic rewiring around glutathione that GSEA failed to highlight.

Finally, scMetNet also identified changes in metabolism around key epigenetic drivers between E13.5 and E17.5, which could be associated with the switch from making neurons to making glia and/or the switch to a quiescent postnatal neural stem cell state. These included genes associated with both Acetyl CoA production and methylation including downregulation of glycolysis associated genes such as *Ldha* and *Hk2*, and downregulation of methionine and SAM metabolism associated genes such as *Dnmt3a, Mat2a*, and *Ahcy*. Upregulation of *Pdha1* in conjunction with downregulation of glycolysis genes suggests diversion of Pyruvate to Acetyl CoA and hence increased OXPHOS. Consistent with the transition from a rapidly-dividing to slowly-proliferating/quiescent state, scMetNet also defined a downregulation of biosynthetic pathways such as PPP and nucleotide synthesis as signified by downregulation of enzymes like *Tkt, Tpi, Prps1* (PPP) and *Paics*.

### Is metabolic rewiring consistent across different regions of the developing brain?

As discussed above we hypothesized that the changes in E17.5 cortical RPs might be part of their transition to a slowly-proliferating/quiescent state. To directly test this idea, and to ask whether this might be a common set of metabolic changes for stem cells, we compared the changes seen in E17.5 cortical RPs to E16.5 dentate gyrus RPs using previously published scRNA-seq data (Hochgerner, Zeisel, Lönnerberg, & Linnarsson, 2018). We found several striking similarities between the two regions (metabolic network shown in Fig. 5). Importantly, both the genes encoding metabolic enzymes and reactions common across the two regions were spread across the whole metabolic map; the common reactions are identified by dashed lines in Fig. 5, including the characteristic upregulation of fatty acid metabolism and the upregulation of mevalonate biosynthesis. Overall, we see thirteen out of the seventeen genes were present in the same metabolic modules. A notable difference is upregulation of Sam metabolism in dentate gyrus, which may partly be due to dentate gyrus data being from E16.5, as Sam upregulation was seen at E15.5 in cortical RPs.

**Figure 5:**
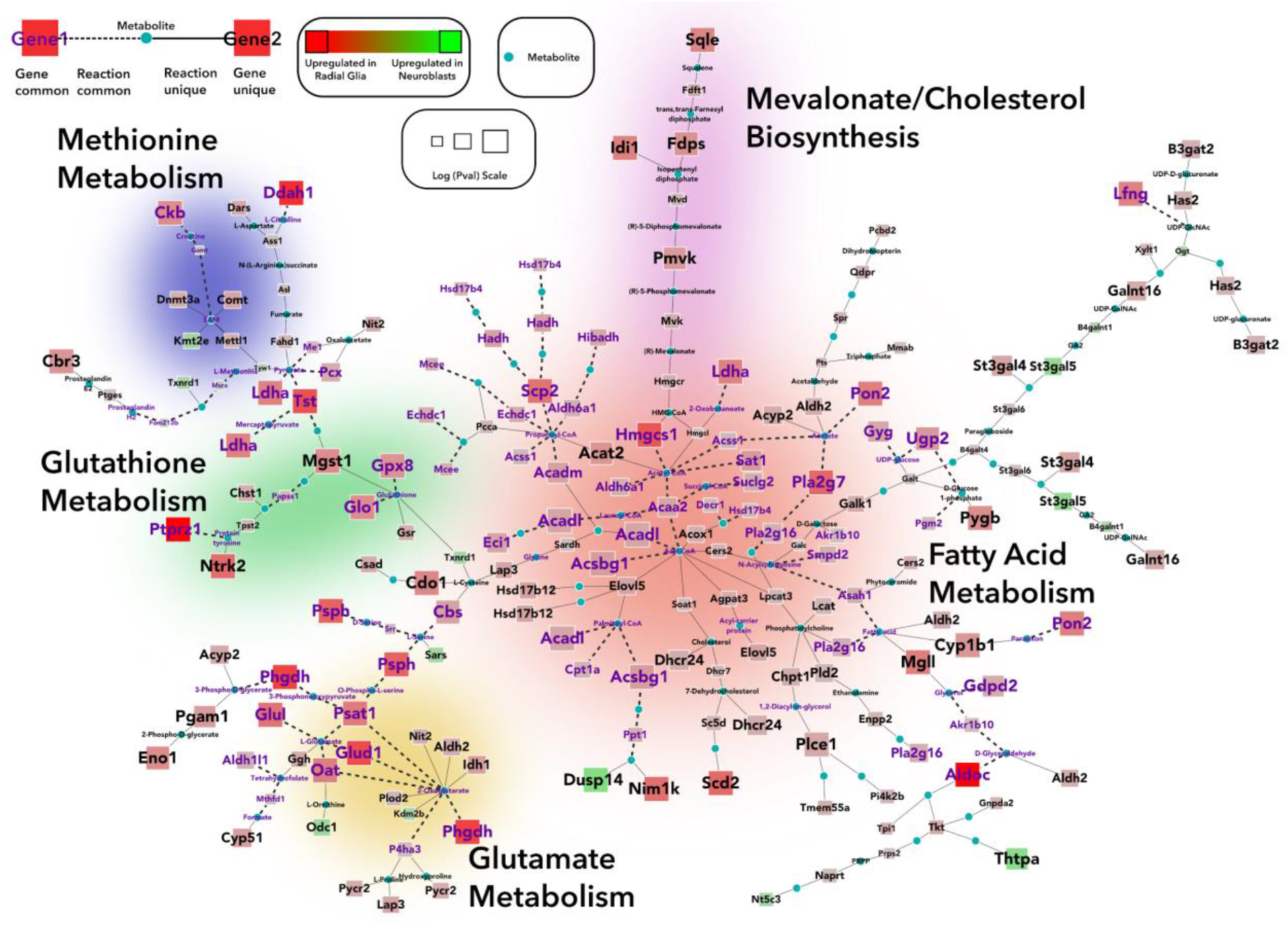
scMetNet network between RPs and Neuroblasts from dentate gyrus at E16.5; the circular nodes represent absent metabolites, square nodes represent genes, increased size of square denotes higher significance. The purple color text denotes that the gene is common between the dentate gyrus comparison and SVZ at E17.5, and the dashed lines represent common reactions.

## Discussion

Single cell transcriptomics has fueled much of the recent rapid progress in understanding cell fate changes during development and tissue regeneration. Here we present scMetNet, a technique that leverages single-cell RNA sequencing data to infer global metabolic rewiring across emerging cell types expected in early developmental events. We first established the congruence between the cell states and metabolic states, both of which are identified using established clustering techniques developed to analyze single cell RNAseq data. To exploit the concordance of metabolic states with cell types we then applied a network-based analysis tool to infer metabolic wiring during early neurogenesis. Our analysis revealed important metabolic modules across the three important functional categories: *bioenergetics, epigenetics, and biosynthesis*. These modules went beyond the results from global analyses such as GSEA that stop at identifying differential pathways, to further pinpoint the underlying genetic and metabolic drivers of the epigenetic machinery responsible for maintaining pluripotency of stem-like cells (Fawal et al., 2018; Shiraki et al., 2014), the rate at which they differentiate, and of the bioenergetic pathways responsible for survival and proliferation this pool of stem-like cells (Hu et al., 2016). Many of these have been suggested in different tissue contexts but have not been fully appreciated and tested in the context of embryonic neural development. One of the surprising findings our analysis uncovered was the upregulation of mevalonate and cholesterol and associated pathways in neural stem cells by stage E17.5. This coincides with a slowing down of RP-mediated neurogenesis. Other shifts such as increased OXPHOS and ROS also point towards the possibility of neural stem cells becoming more quiescent or dormant. High levels of cholesterol biosynthesis have been previously reported during neural development and in adult brains astrocytes/microglia. However, the role of cholesterol production by neural stem cells as their neuronal output slows down at E17.5 remains unclear. Previous work where alterations in cholesterol synthesis in stem cells led to neuron apoptosis rather than impaired neurogenesis (Saito et al., 2009) suggests a possible cell extrinsic role of cholesterol produced by the neural stem cells.

We would like to reiterate the subtle “signals” in metabolic rewiring, such as increased ROS activity at E17.5 and the associated genes and metabolites driving the increase remain hidden in global approaches such as GSEA. This is not surprising as GSEA relies on the a priori definitions of pathways, where each pathway is exhaustively defined across different biological contexts and hence have many more genes and metabolites that play out in any particular biological context. The output from the scMetNet analysis, on the other hand, is independent of such global pathway definitions and aims to build a network of connected metabolic reactions independent of the pathway they are associated with. Hence, scMetNet facilitates painting a holistic picture by directly providing interactions between the different pathways.

Now, armed with a detailed knowledge of metabolic rewiring at the level of individual genes and metabolites we explore here potential interventions to alter embryonic neural development with focus on epigenetic and bioenergetic modules. Our analysis identified twelve genes associated with epigenetic metabolites, acetyl-CoA and a-KG, one-carbon metabolism and methylation-demethylation associated genes. Out of the twelve, seven genes have been implicated in neuropathologies but have not yet been directly explored in their role in altering early neural development. For instance, deficiency of the two a-KG associated enzymes *Psat1* and *Phgdh*, which are upregulated in RPs, have been reported in microcephaly, psychomotor retardation and epilepsy (Acuna-Hidalgo et al., 2014; Sharma & Prasad, 2017; Yoshida et al., 2003). We hypothesize that a KO of *Phgdh/Psat1* will advance RP differentiation timing to an earlier developmental stage. In the same vain, we hypothesize that the perturbation of mitochondrial genes associated with acetyl-CoA (*Acadl, Acadm, Acss1, Hadh, Acaa2*) will lead to a smaller pool of RPs due to decreased energy production, while perturbing acetylation via *Hat1* will lead to early differentiation of RPs.

The coincidence of slowing down of RPs neural output at E17.5 and upregulation of mevalonate and cholesterol provide a set of potential targets for altering the RPs neuronal output. Our anlysis suggets application of statins - inhibitors of *Hmgcr* - will promote neurogenesis. Previous work suggests this modulation may act through Wnt signaling (Robin et al., 2014), consistent with which we found upregulation of *Lfng* which is a part of Wnt signaling pathway in RPs at E17.5. Similarly, deficiency of *Dhcr7*, important for cholesterol synthesis, is known to cause Smith-Lemli-Opitz syndrome (Liu et al., 2014), and we hypothesize perturbing its levels will alter the neuronal output in embryos.

More broadly, many of the genes that define the metabolic rewiring network have been associated with cancers, which is not surprising considering the connection between stemness and tumorigenesis. The genes *Psat1, Phgdh* and *Ptprz1* are known to be upregulated in glioblastoma and lower grade glioma. Both *Psat1* and *Phgdh*, which catalyze important biosynthetic reactions, have been extensively studied in many cancers (Amelio et al., 2014), and *Ptprz1* has been said to inhibit stem cell like properties of tumor cells (Fujikawa et al., 2017). Here we have highlighted a few examples of potential perturbations to alter embryonic neural development, please see the details for all the genes we find in the rewiring network and their potential effect in the Supplementary Table S4 and highlighted in Supplementary Fig S7.

Overall, in this paper we show that scMetNet can utilize single cell data to both identify key metabolic pathways and then go further and identify specific genes, metabolites, and reactions that drive those metabolic pathways. Here by applying our approach to early neural development we propose a specific set of perturbations that can be used to alter different aspects of neurogenesis. Our approach is general and can be carried over to single cell data from other tissues either during development or regeneration to discover novel therapeutic targets as discussed above in the context of neuropathologies.

## Materials and Methods

The scMetNet pipeline combines single cell analysis (Butler, Hoffman, Smibert, Papalexi, & Satija, 2018; Stuart et al., 2019) with network analysis (Jha et al., 2015) into an integrated pipeline that can take single cell data and perform all the analysis detailed below. The pipeline is available to run with the data used in the paper at Polly (https://polly.elucidata.io/).

### Data Processing and Differentially Expressed Genes Selection

Gene expression matrices of cortical cells at all three embryonic ages (E13.5, E15.5, E17.5) were used from the GEO accession: GSE107122. All of the analyses and plots in the paper have been done using R (version 3.5.2) and Seurat (V2) package (Butler et al., 2018). We used the following quality control steps to reproduce the analysis done by Yuzwa et al: (i) Cells that expressed less than 200 genes and cells that expressed more than 2500 genes were excluded from the analysis, lower than 200 genes is an indicator of lower quality while higher than 2500 genes indicates that the cell might be a potential doublet; (ii) Genes expressed by less than 3 cells were removed; (iii) Cells having more than 15% mitochondrial content (as denoted by genes derived from the mitochondrial genome) were removed. The data were normalized using the *NormalizeData* function keeping normalization method as *LogNormalize* and scale factor as 10000. Unwanted variation due to total RNA in cells and mitochondrial gene content were regressed out using the *ScaleData* function in Seurat. Highly variable genes were computed for the counts at each embryonic age using the *FindVariableGenes* method of Seurat, we used *ExpMean* as the mean function, *LogVMR* as the dispersion function, mean lower threshold as 0.0125, mean higher threshold as 8 and dispersion threshold as 0.05. Lower dispersion threshold was used to get a higher number of genes as scMetNet needs genes that show both high and low variability to ascertain the background signal and identify significant and connected subnetworks. Using this method, we got 5888, 6539 and 5265 highly variable genes for the embryonic ages E13.5, E15.5, E17.5 respectively.

PCA was done using the highly variable genes and the first 20 principal components for each embryonic age were used for clustering and TSNE implementation. Clustering was performed using the SNN-Cliq method implemented in Seurat with a resolution of 2.5 for all three embryonic ages. To visualize clusters, t-SNE projections were calculated on the principal components using the default perplexity of 30. Annotation of clusters was done using the expression of known marker genes for each population as mentioned in Supplementary Table S1.

Differential expression analysis was done between cell population clusters using the Wilcoxon Rank Sum test available in Seurat.

### Combine Single-Cell Datasets

Gene expression matrices from E13.5 and E17.5 were combined using Canonical Correlation Analysis available in Seurat. A union of highly variable genes from the counts at the two embryonic ages was used and the number of canonical vectors to calculate was set to 20.

### Metabolic state extraction

For finding metabolic states, the dataset was limited to the 5266 metabolic genes identified from KEGG before processing. Rest of the steps followed were identical to the procedure described above except at the clustering step the resolution parameter was chosen as 0.5, 0.3, 0.4 at the three embryonic ages E13.5, E15.5 and E17.5 respectively to get three clusters corresponding to the three metabolic states at E13.5, E15.5 and two clusters corresponding to two metabolic states at E17.5. For figure S1C, a set of 4000 random genes was sampled from all genes and the pipeline mentioned above was run for each random gene set. The random gene set sampling was done 1300 times. The overlap between cells for each cell type and corresponding metabolic state was found for each random gene set and compared to overlaps obtained for states obtained using only metabolic genes. The p-values for the cell identities assigned using metabolic genes clustering were calculated using the permutation test including all the permutations.

### Network Analysis

Network analysis pipeline prescribed by Sergushichev et al (Sergushichev et al., 2016) was used to construct the maps depicted in all figures. The network analysis pipeline performs the integrated network analysis of transcriptional and metabolomic data to find the most changing subnetworks in the KEGG database. The pipeline was run using Intomix which is a Polly (https://polly.elucidata.io/) software.

Intomix uses a similar pipeline as mentioned in Sergushichev et al. We used the latest kegg database as of 2019 (https://www.genome.jp/kegg/) to construct the global network using KEGG REACTION database. We downloaded enzymes, glycans, compounds and reactions from KEGG and made a global network by removing ubiquitous metabolites such as ATP and collapsing groups of anomeric metabolites into one metabolite. KEGG ENZYME database was used to make the enzyme-gene-organism mapping.

The analysis was performed as a two-step process -

1. Creating a network of reactions based on the scRNA differential expression data - The algorithm converts the differential expression of genes into the differential expression of reactions. Genes taking part in a specific reaction and coding for an enzyme are taken into account and the minimum pval represented within the gene set is selected as the p-value of the reaction. Reactions with no assigned p-values are dropped. The reactions are interpreted as a node for the purpose of network creation.
2. Finding a reaction module - After creating an optimal network, a connected reaction module representing significant changes between the cell clusters is identified. The optimal reaction module is found out based on the scoring of nodes and edges as per the assigned p-values. All modules are found using the Heinz solver, keeping the solving time to 4 minutes.

### Metabolic genes

The set of metabolic genes used for analysis in Figure 1 was obtained from the KEGG pathway database.

### Gene Set Enrichment Analysis

Gene Set Enrichment Analysis (GSEA) was done at each embryonic age using gene sets of metabolic pathways which were obtained from the KEGG database. The R package fgsea (Sergushichev, 2016) was used to perform this analysis. GSEA was done on pathways from the KEGG database which had more than 5 genes and less than 500 genes and the number of permutations to run was set to 100,000. All the GSEA results can be found in Supplementary Table S2 and S3.

### Pathway over-representation analysis

Pathway over-representation analysis was done using the tool Enrichr (Kuleshov et al., 2016). All the genes picked by scMetNet were used to make a heatmap. The average expression of these genes for Neurons and RPs at the three embryonic stages were clustered according to embryonic days. Gene sets corresponding to each cluster were then used to find over-represented pathways among Neurons and RPs. The heatmap thus obtained is shown in Supplementary Figure S6.

## Code Availability

Complete code for the analysis done in this paper can be found at: https://github.com/ElucidataInc/single_cell_metabolism_scripts

## Acknowledgements

We would like to thank Richard Kibbey for critical readings of the manuscript and Gary Bader for many discussions. SG and FDM were supported by funding from the CFREF “Medicine by Design” program. FDM. is a Canada Research Chair and an HHMI Senior International Research Scholar.

## Author contributions

SG and AJ conceived the study. SJ and SM analyzed data and implemented the scMetNet pipeline. SJ, SM, SG and FDM cowrote the manuscript.

## Supplementary figures

**Figure S1:**
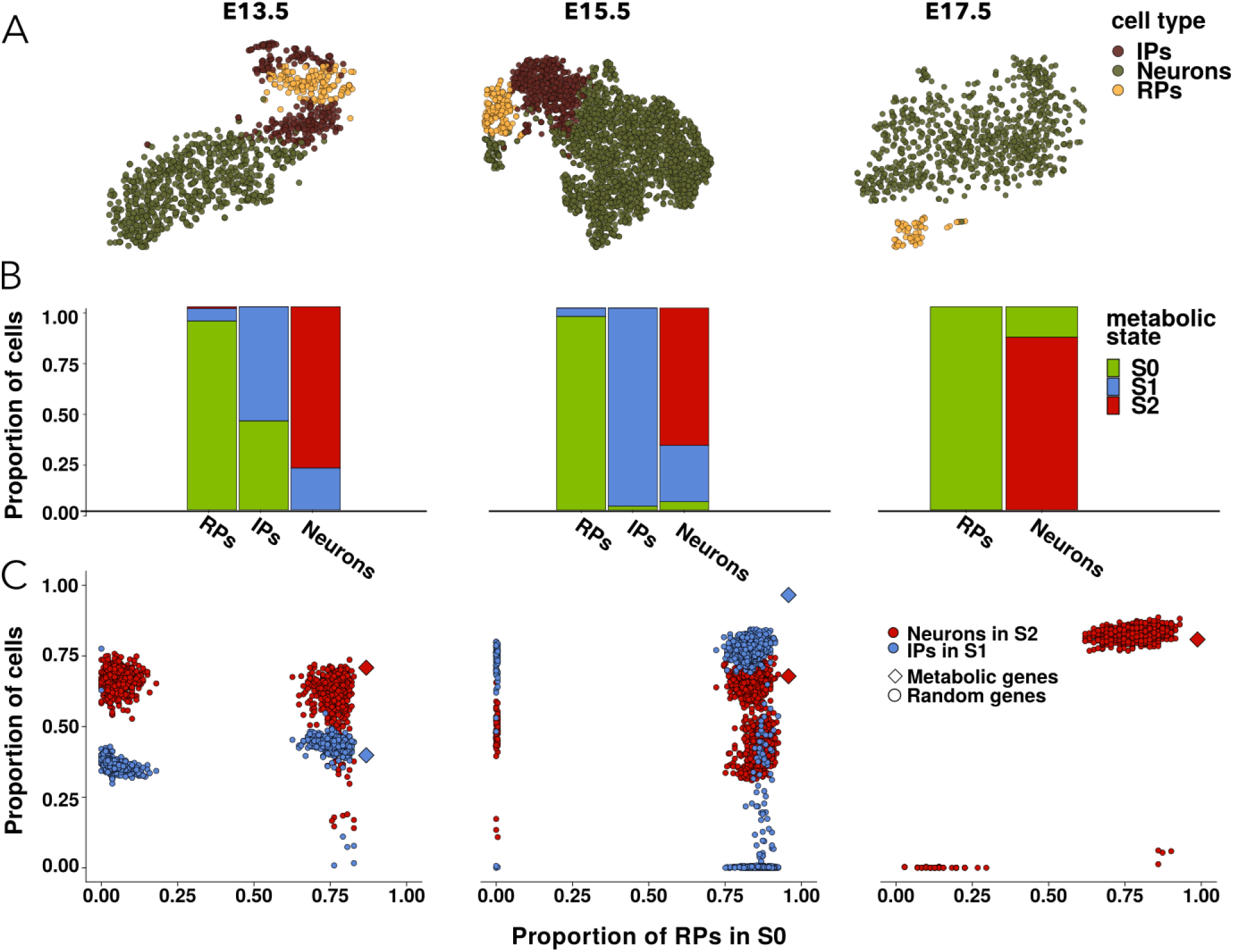
(A) t-SNE visualization of cell clusters at E13.5, E15.5 and E17.5 using all the variable genes. (B) The overlap of cell types and metabolic states at E13.5, E15.5 and E17.5. (C) Cells were clustered using random sets of genes, the proportion of RPs in metabolic state S_0_ (green), IPs in metabolic state S_1_ (blue) and Neurons in metabolic state S_2_ (red) were calculated. The p-values for the cell identities assigned using metabolic genes clustering calculated using the permutation test are 7.77e-4, 7.44e-4 and 7.33e-4 for E13.5, E15.5 and E17.5 stages, respectively.

**Figure S2:**
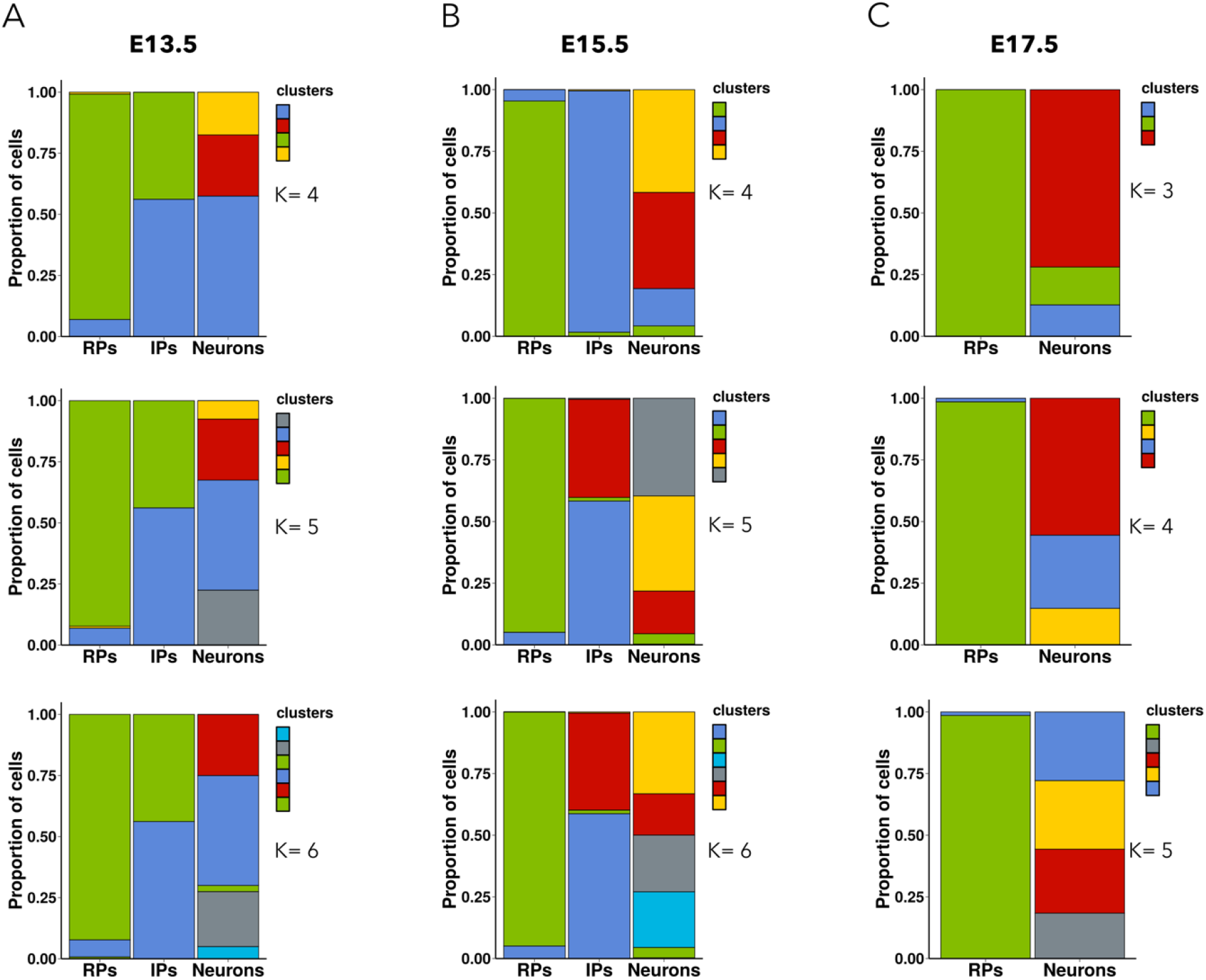
The proportion of cells from the clustering obtained using metabolic genes, present in the three cell type clusters with an increasing number of clusters at (A) E13.5 (B) E15.5 and (C) E17.5.

**Figure S3:**
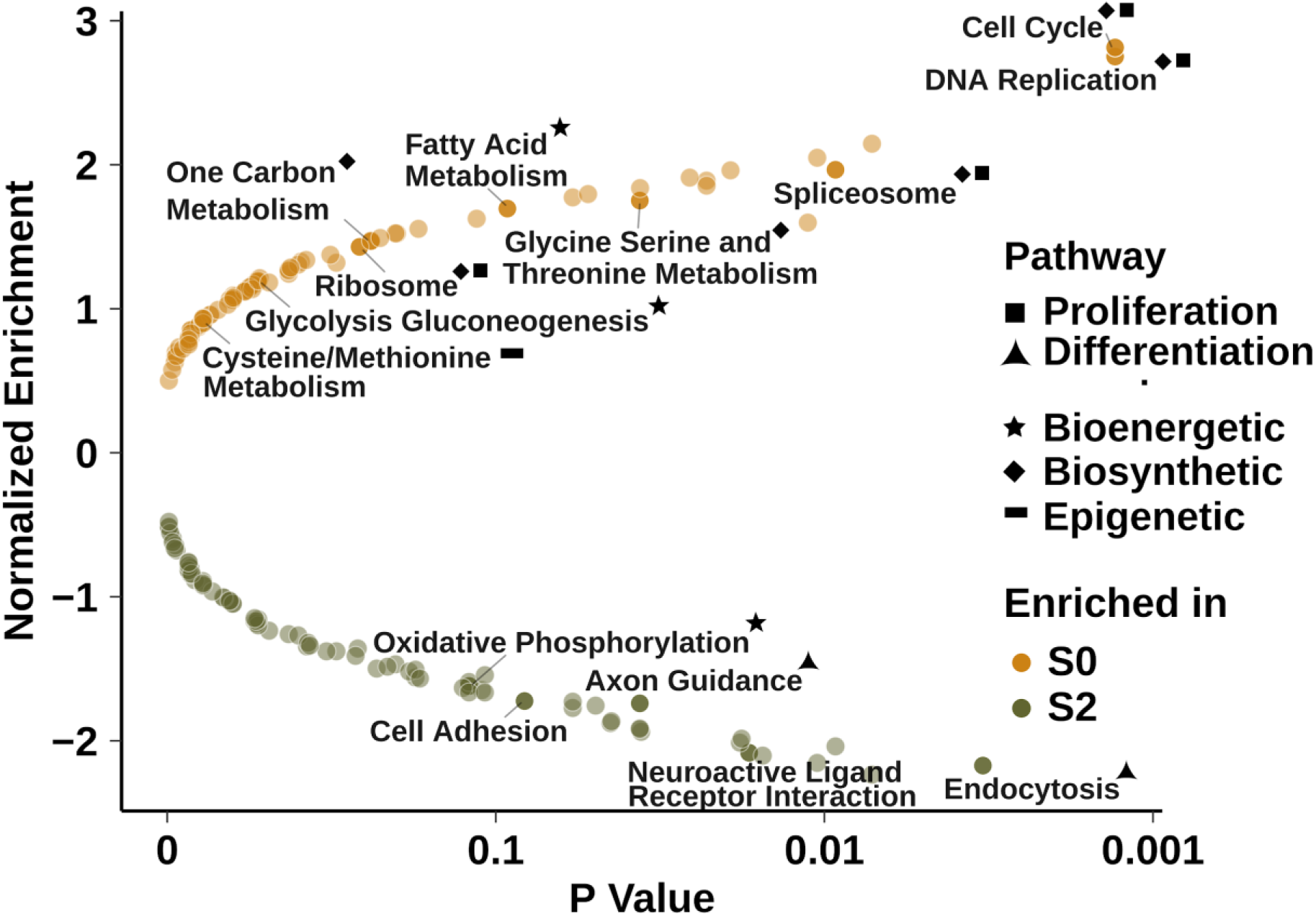
Gene Set Enrichment Analysis (GSEA) between S_0_ and S_2_

**Figure S4:**
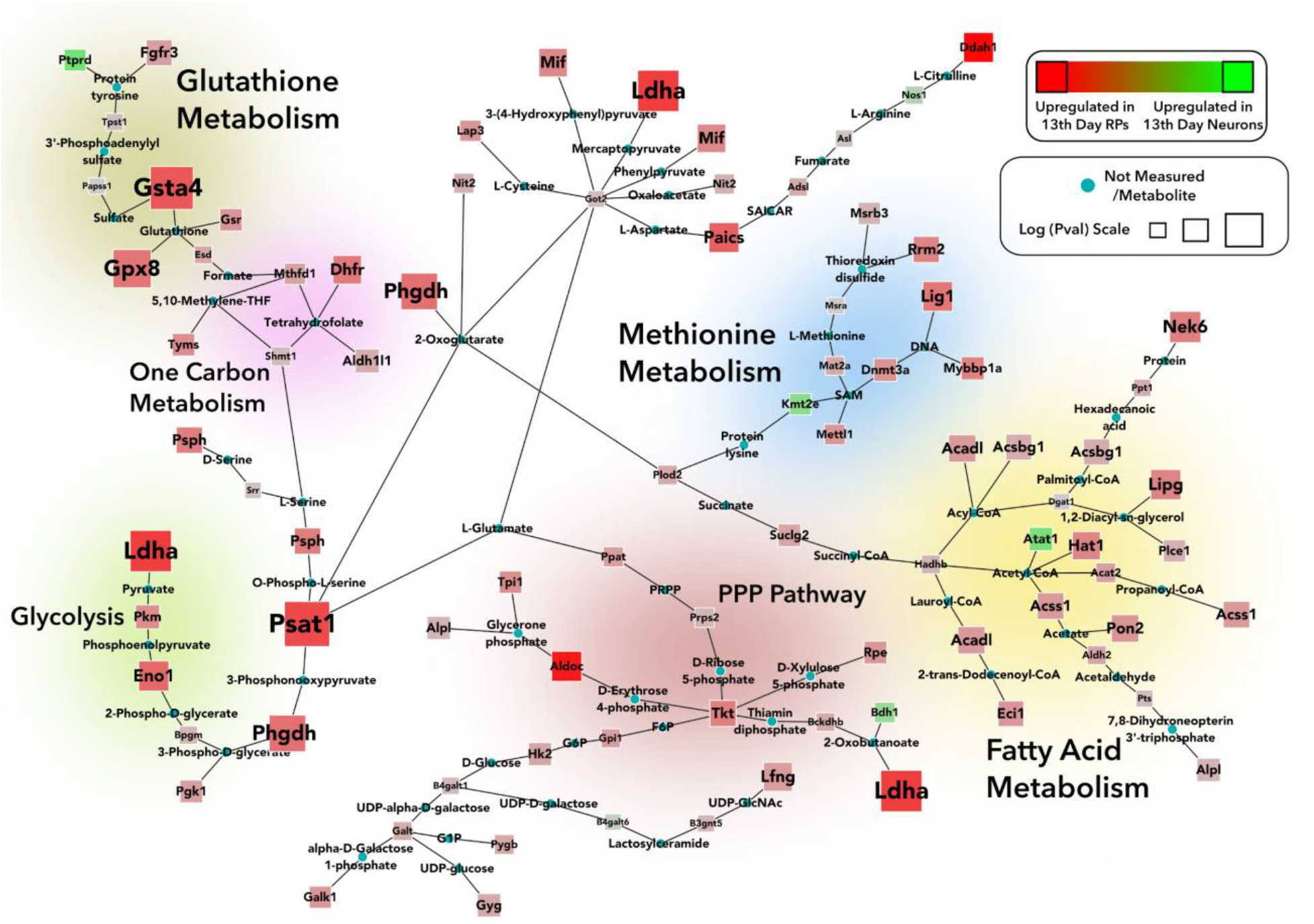
scMetNet network between RPs vs Neurons at E13.5; the circular nodes represent metabolites that connect the metabolic enzymes represented by square nodes; the size of square nodes around the enzymes denotes significance.

**Figure S5:**
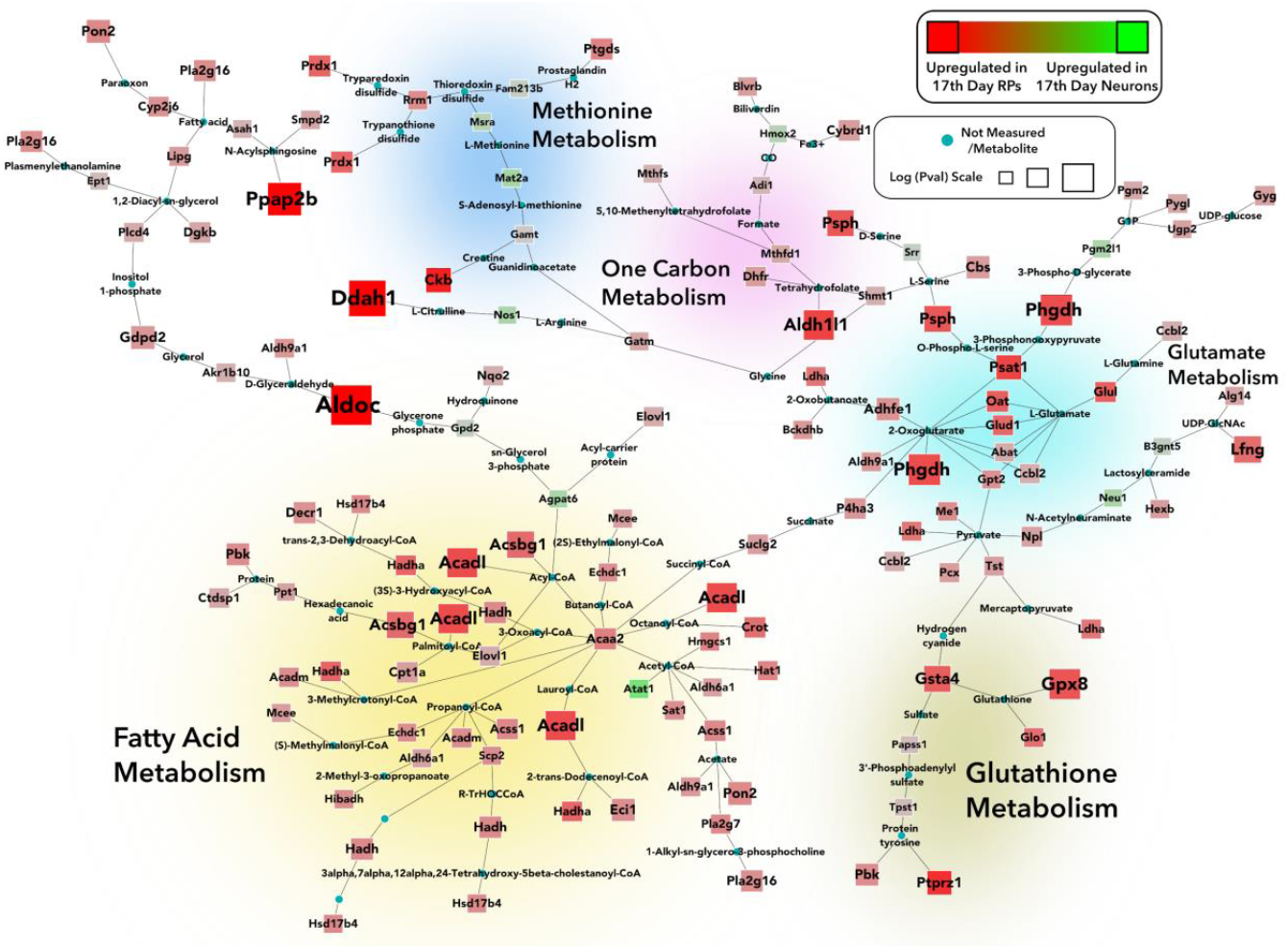
scMetNet network between RPs vs Neurons at E17.5; the circular nodes represent metabolites that connect the metabolic enzymes represented by square nodes; the size of square nodes around the enzymes denotes significance.

**Figure S6:**
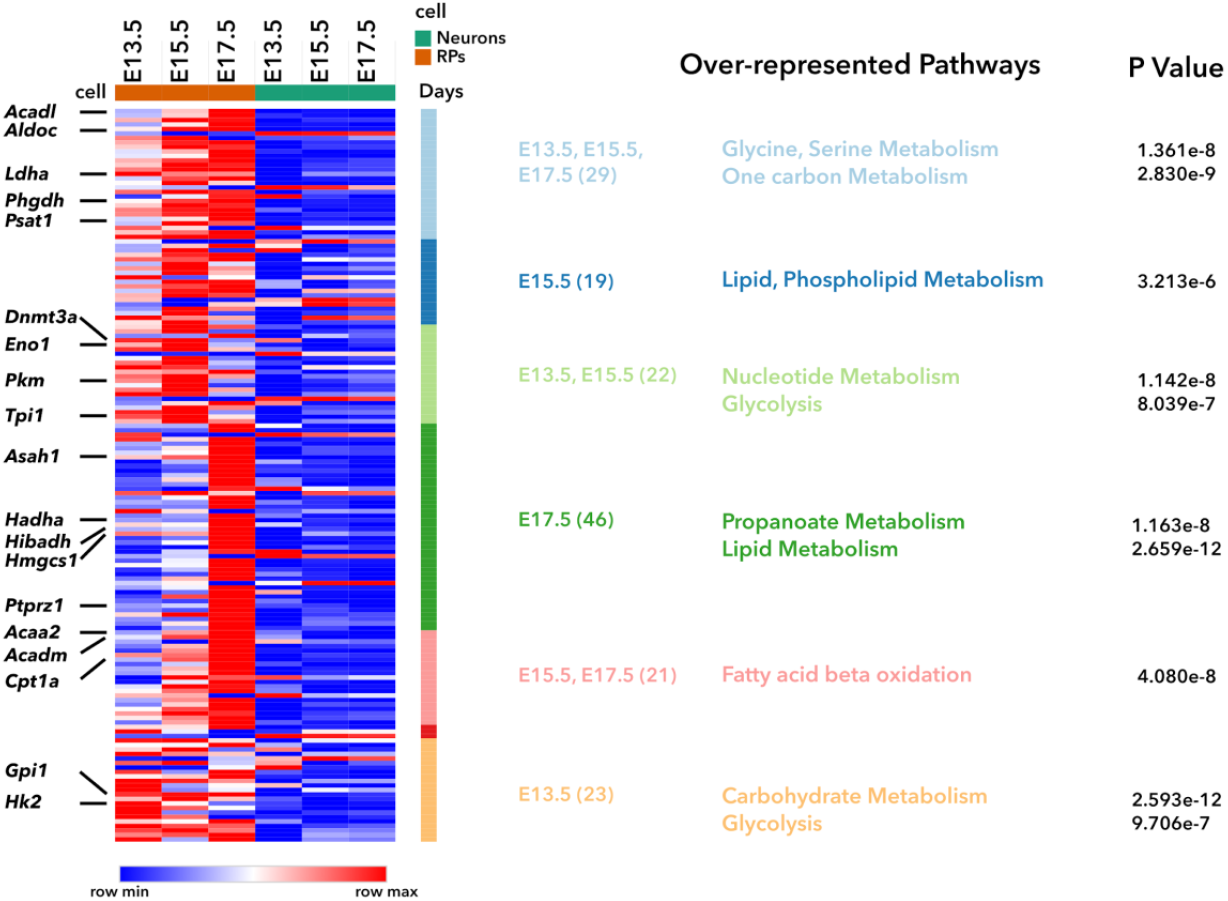
Heatmap of genes picked up by scMetNet for E13.5, E15.5, and E17.5 stages; same genes are shown in the Venn Diagram in main text Figure 3B. Enriched pathways are shown for each stage individually, pair of stages, and all the stages together. The number of genes considered are mentioned in parenthesis. Over-represented pathways for each day clusters along with p values are mentioned (see Methods for details).

**Figure S7:**
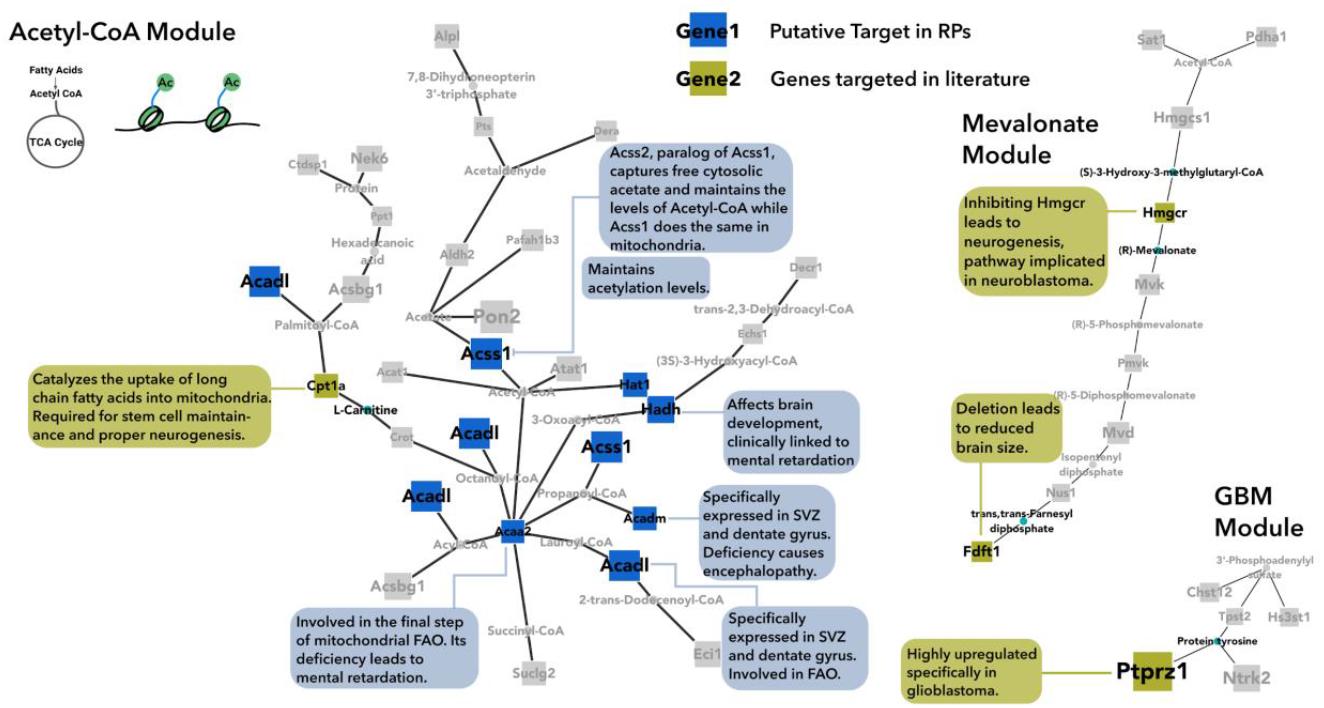
Different modules/metabolites defining the epigenetic and bioenergetic landscape of RPs and their relationship to past metabolic studies, and new potential genetic targets to alter early brain development (also see Supplementary Table S4 for a comprehensive list).

**Table S1:** Marker Genes for the three time points.

| E13.5 |  |  | E15.5 |  |  | E17.5 |  |  |
| --- | --- | --- | --- | --- | --- | --- | --- | --- |
| RP | IP | Neurons | RP | IP | Neurons | RP | IP | Neurons |
| Sox2 | Eomes | Tbr1 | Sox2 | Eomes | Tbr1 | Sox2 | Eomes | Tbr1 |
| Pax6 | Gadd45g | Tubb3 | Pax6 | Gadd45g | Tubb3 | Pax6 | Sstr2 | Tubb3 |
| Hes1 | Neurod1 | Satb2 | Hes1 | Neurod1 | Satb2 | Hes1 |  | Satb2 |
| Hes5 | Ngn1 | Bhlhe22 | Hes5 | Ngn1 | Bhlhe22 | Hes5 |  | Bhlhe22 |
| Slc1a3 | Slc17a6 |  |  | Slc17a6 | Unc5d | Slc1a3 |  |  |
|  | Ngn2 |  |  | Ngn2 | Sema6d |  |  |  |
|  | Btg2 |  |  | Btg2 |  |  |  |  |
|  | Sstr2 |  |  | Sstr2 |  |  |  |  |
|  | Mfap4 |  |  | Mfap4 |  |  |  |  |

**Table S2:**
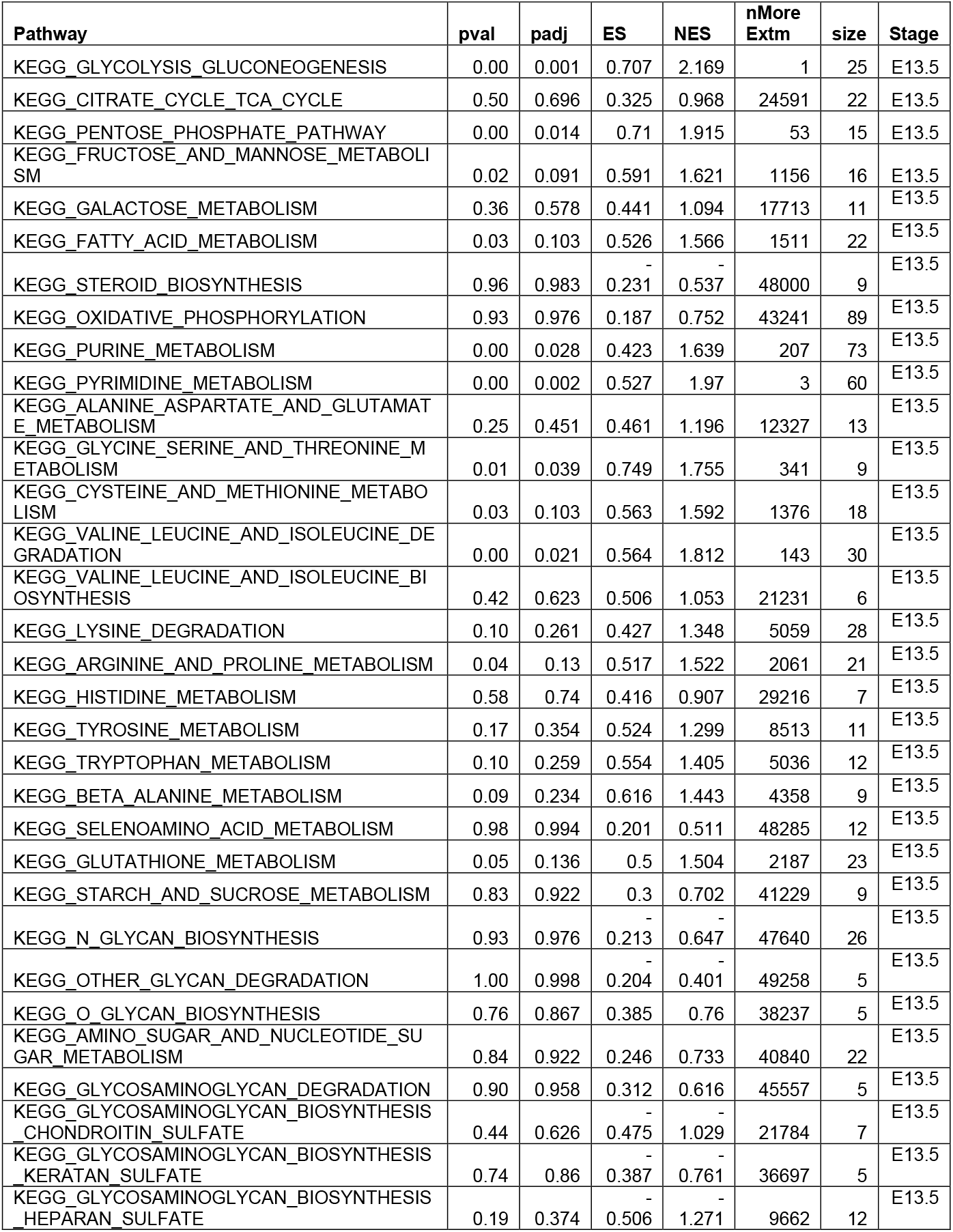
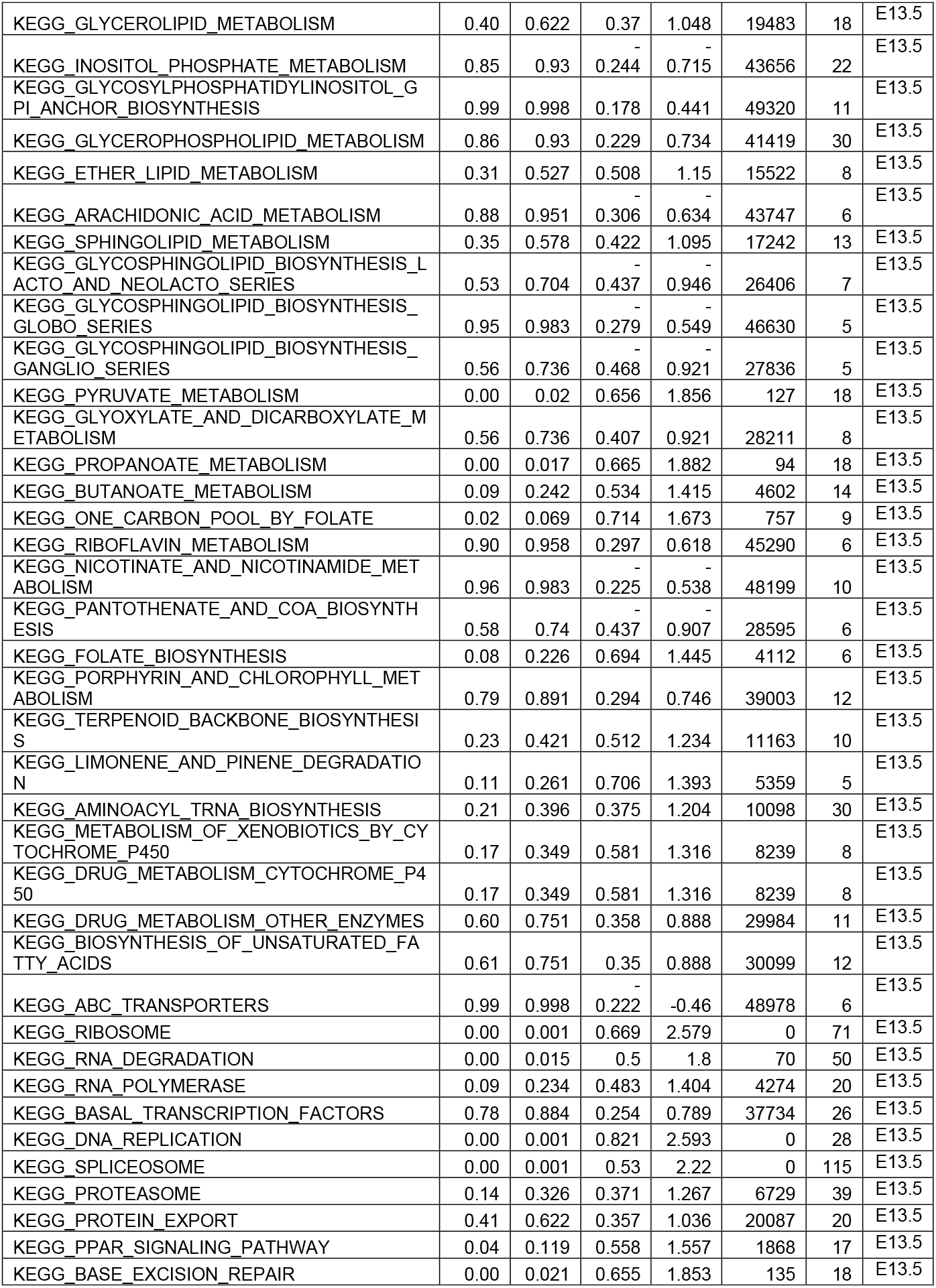
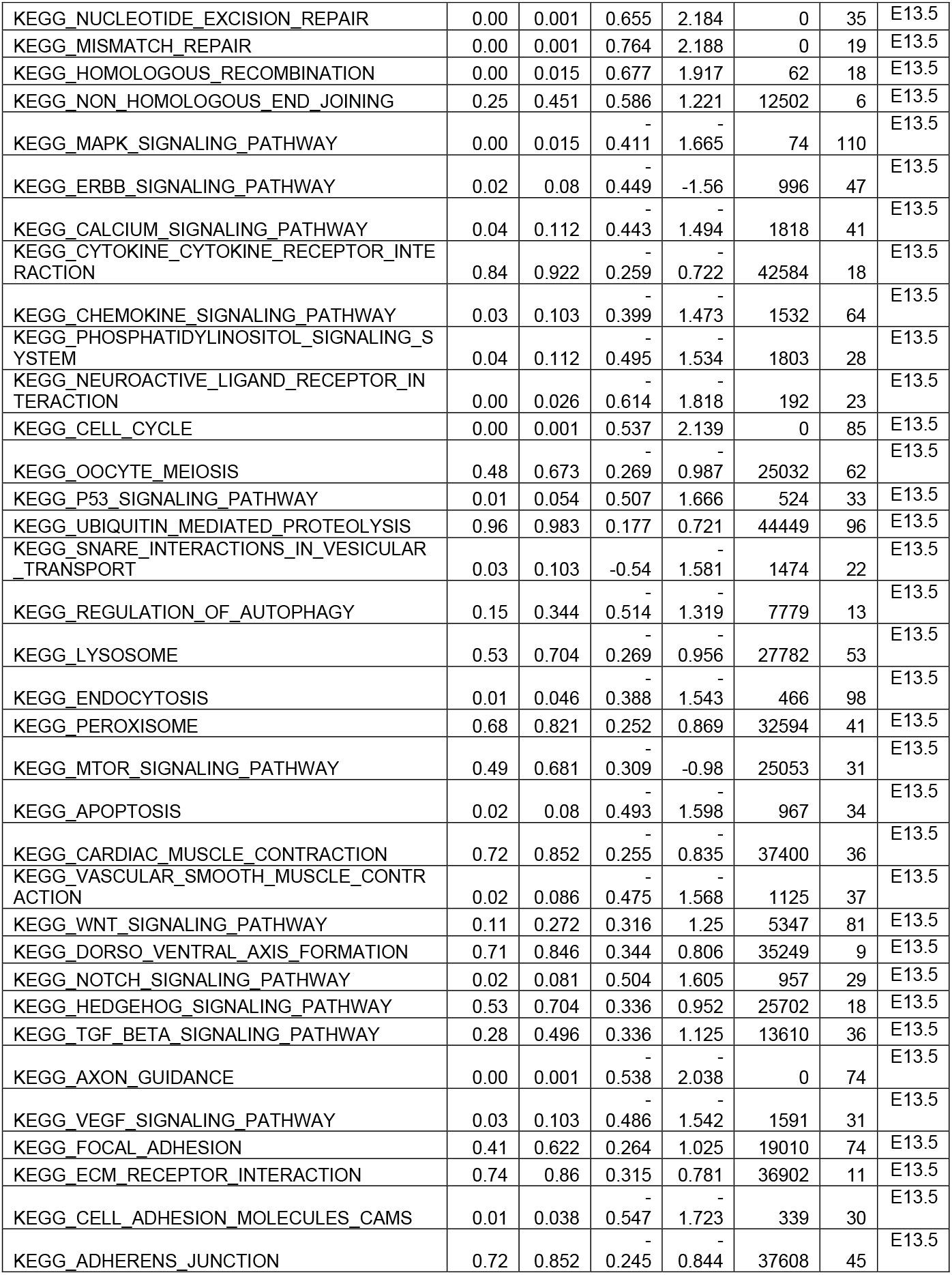
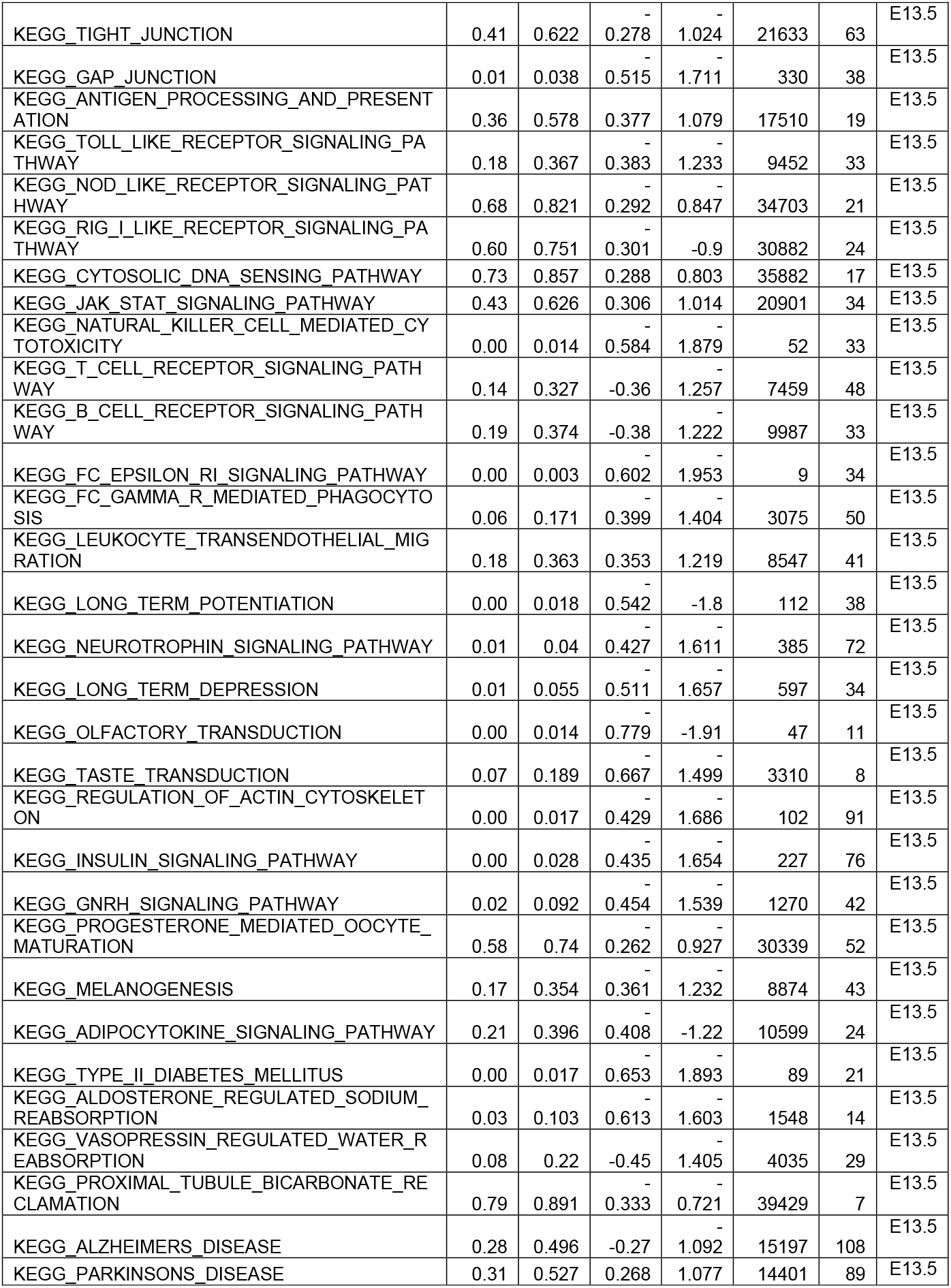
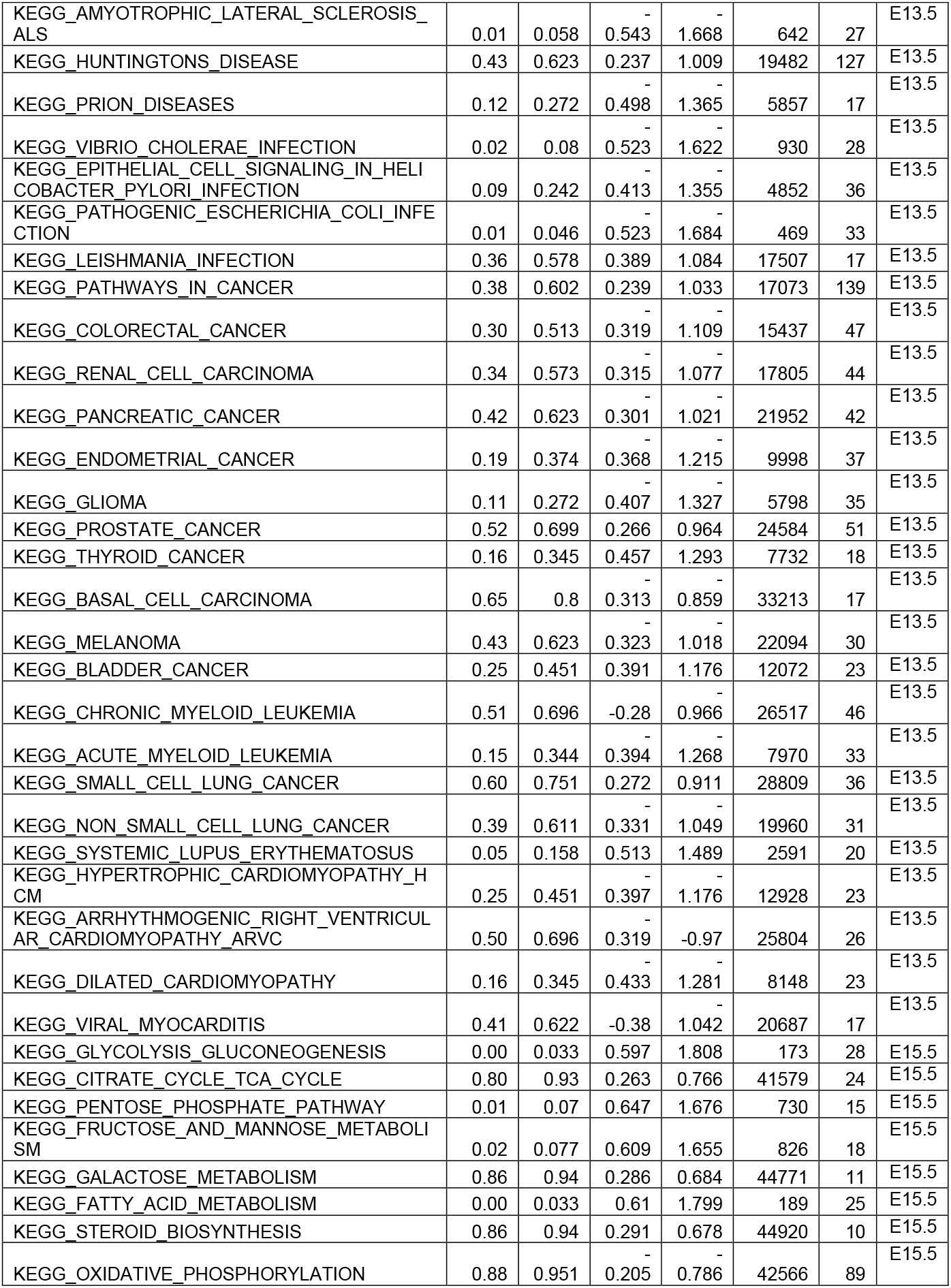
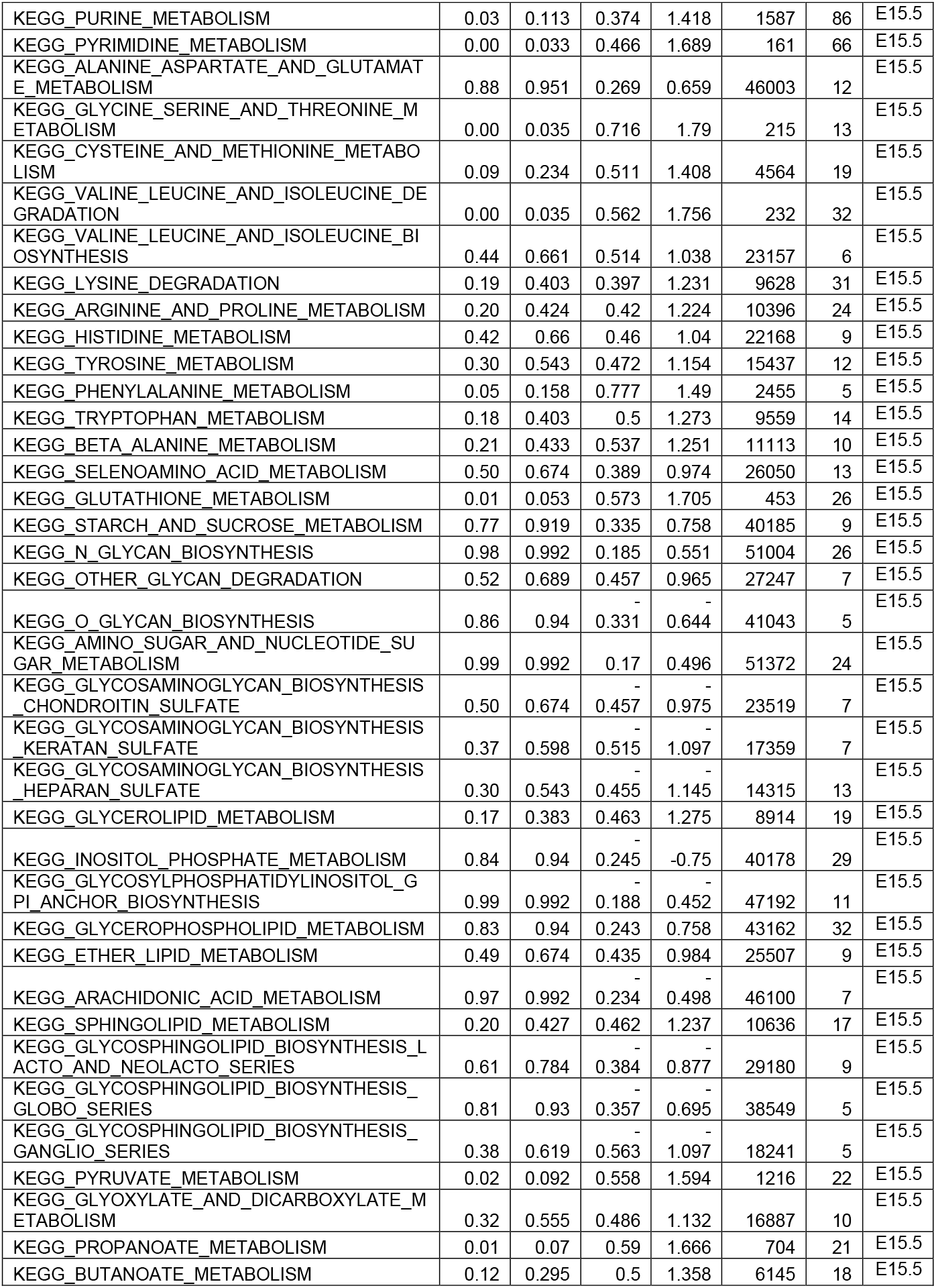
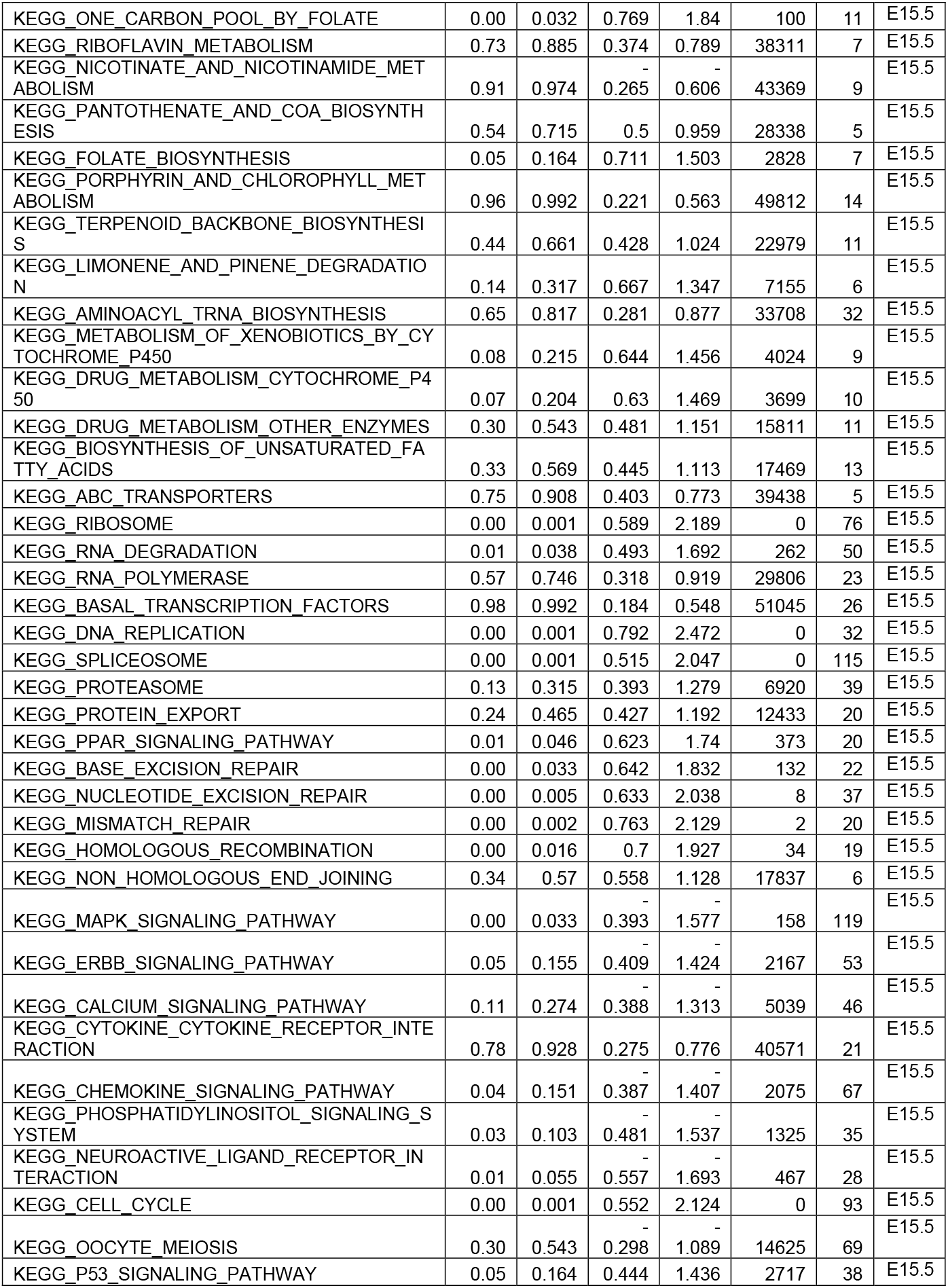
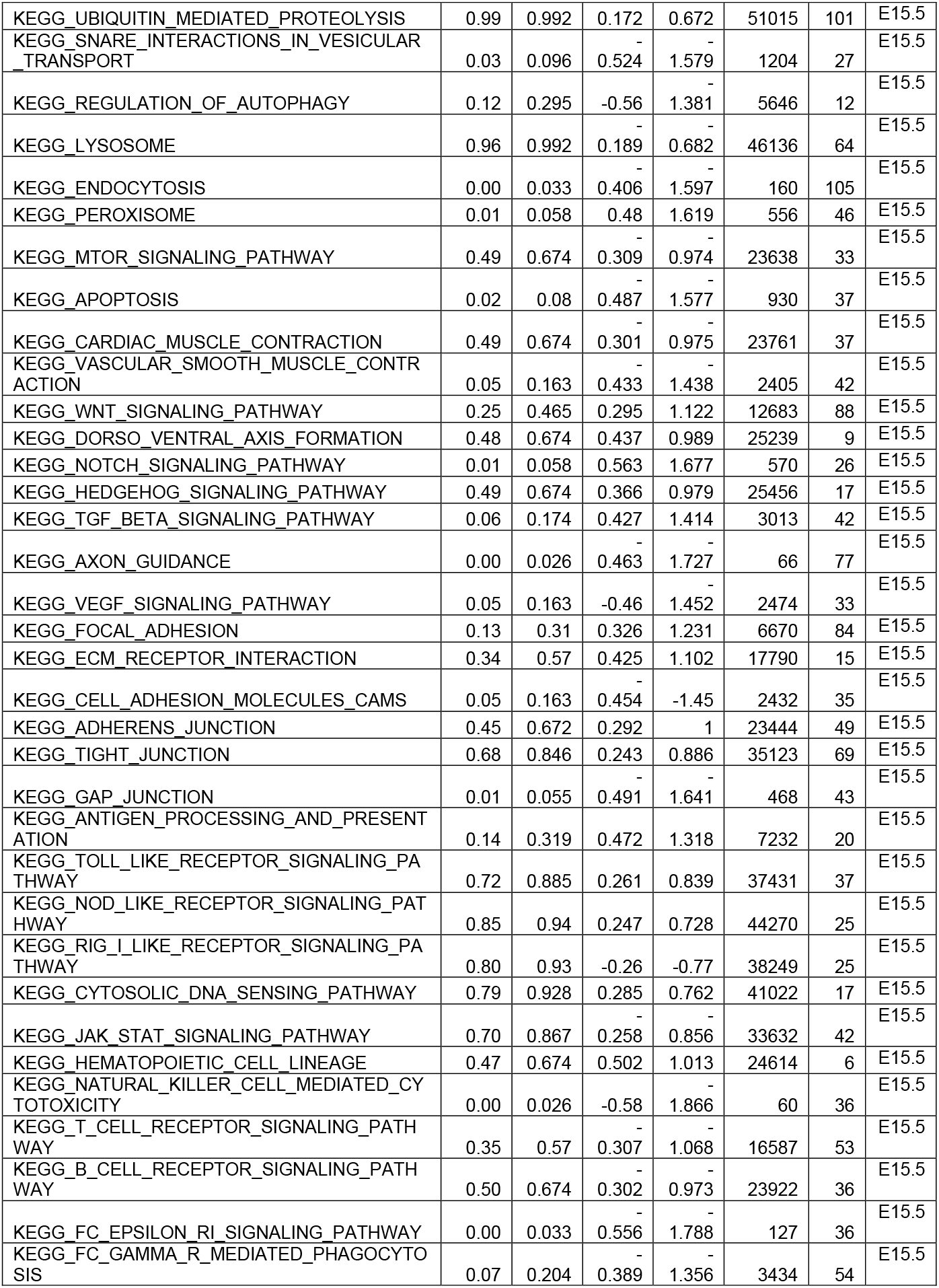
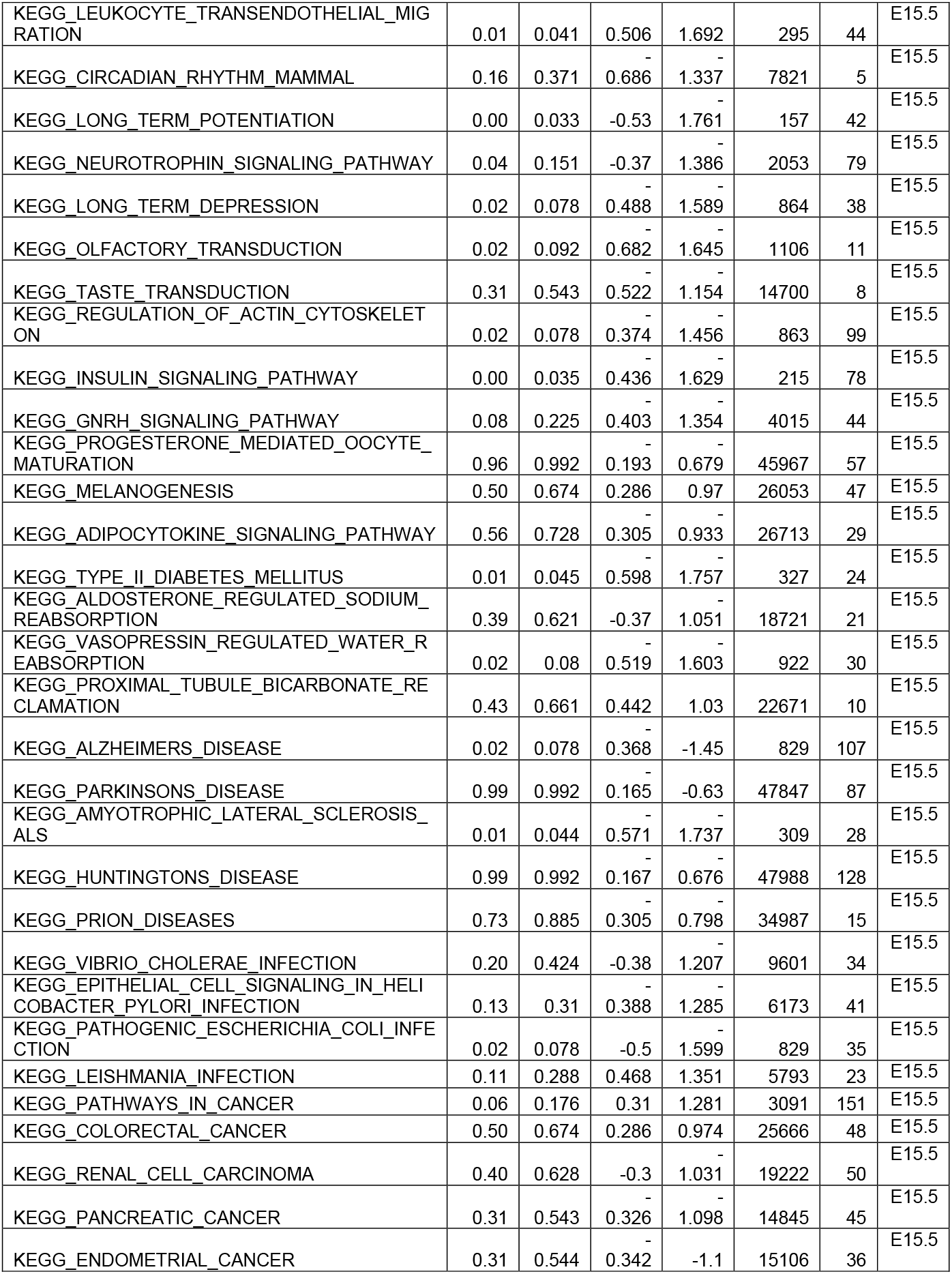
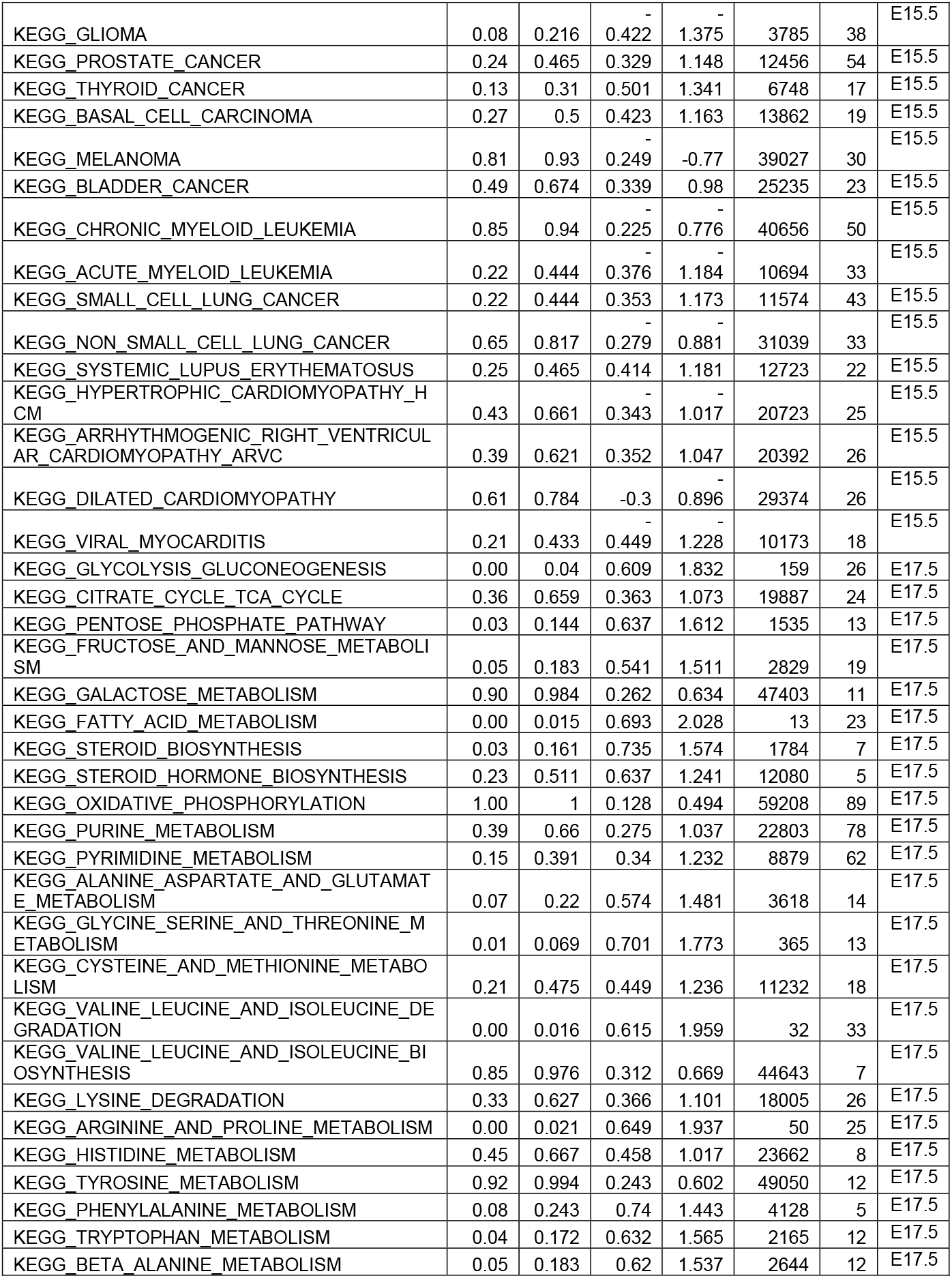
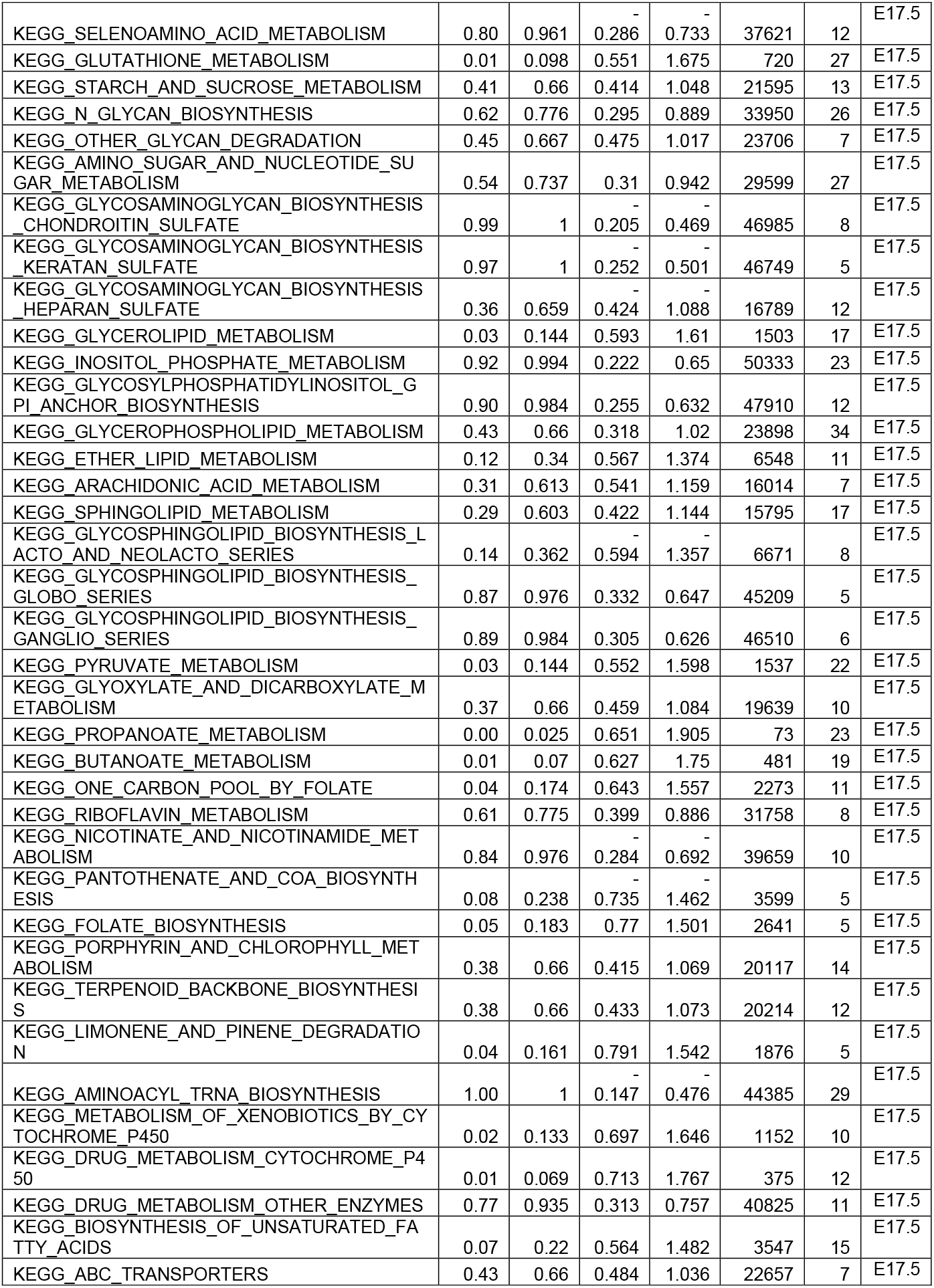
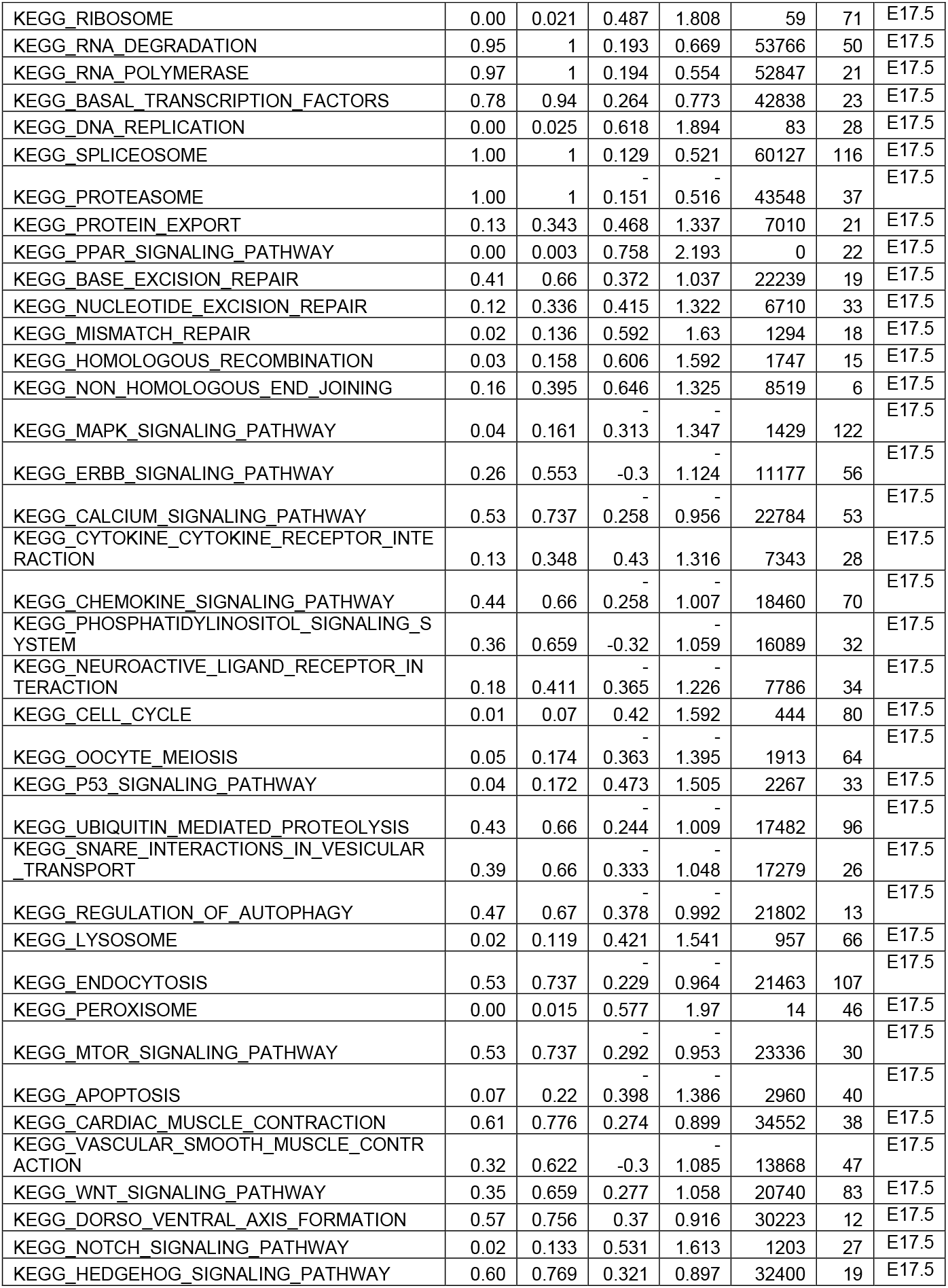
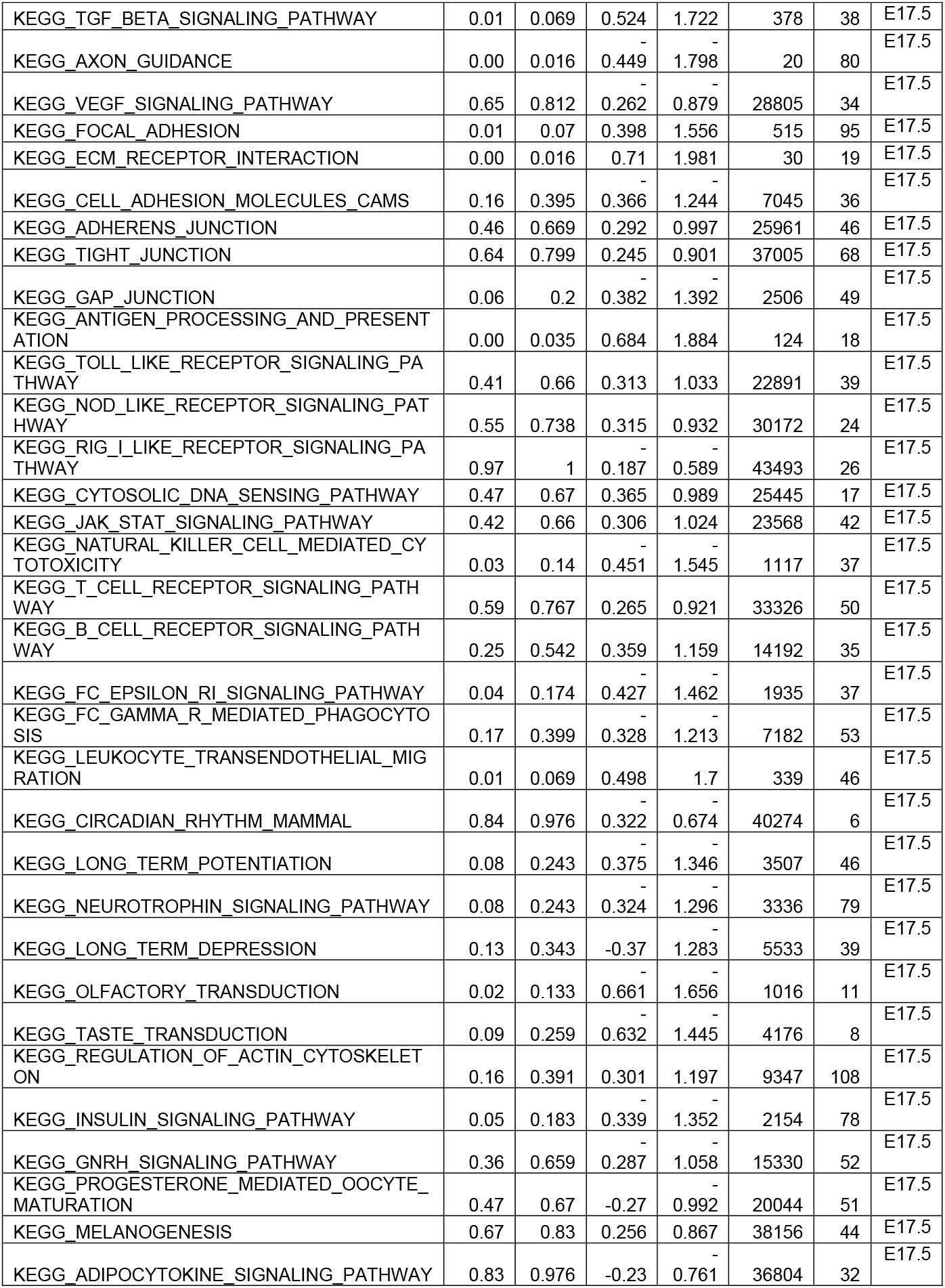
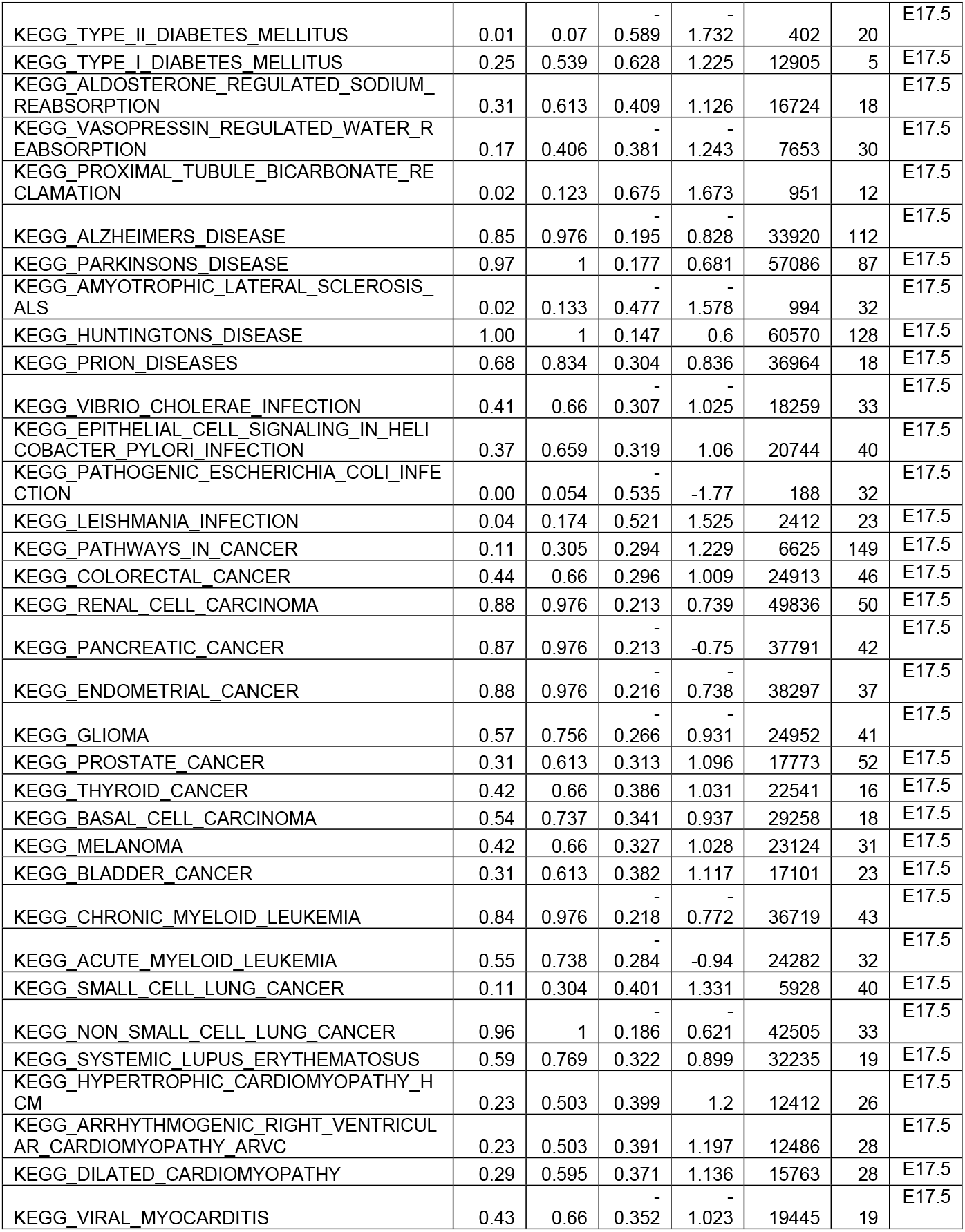
RPs vs Neurons fgsea all days.

**Table S3:**
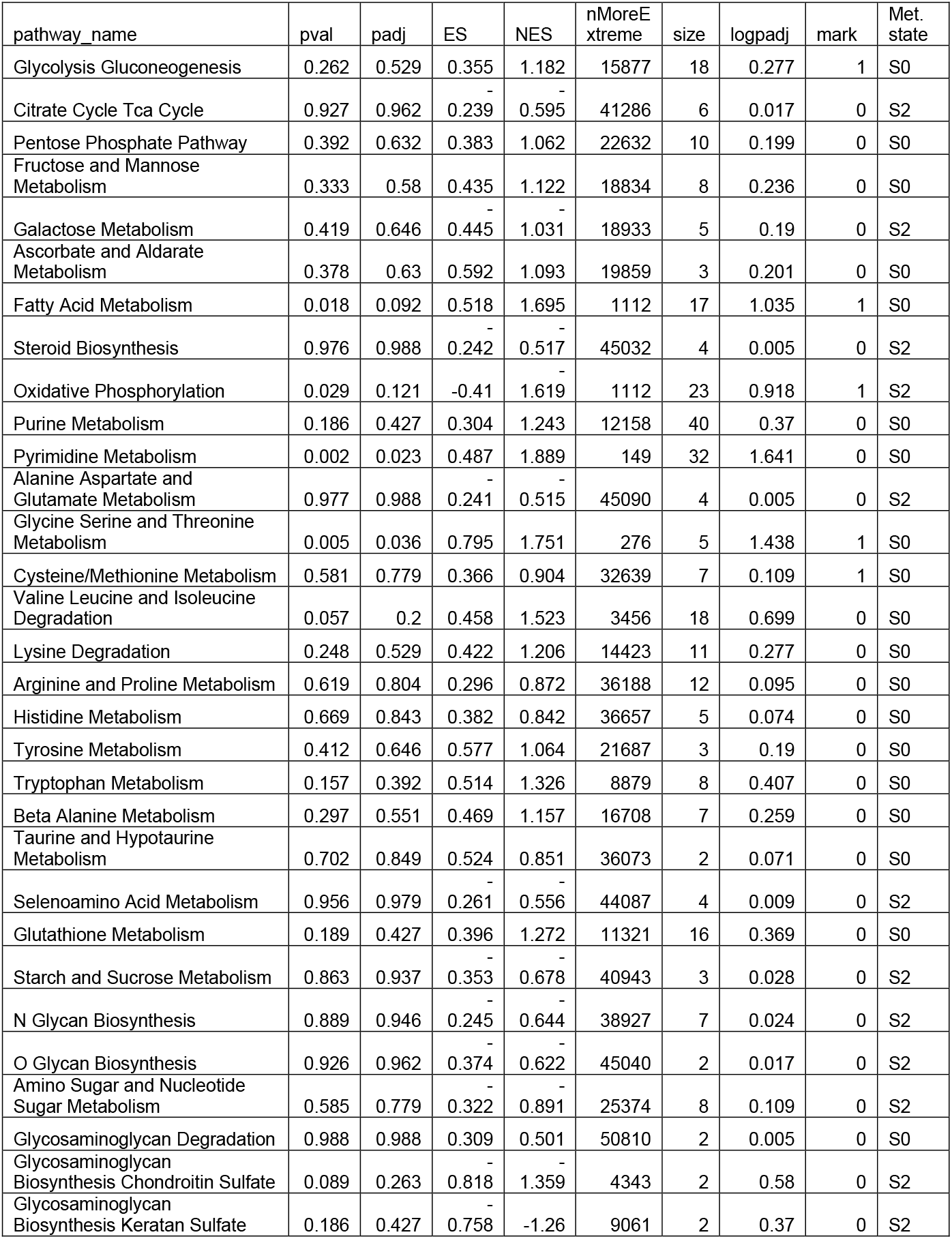
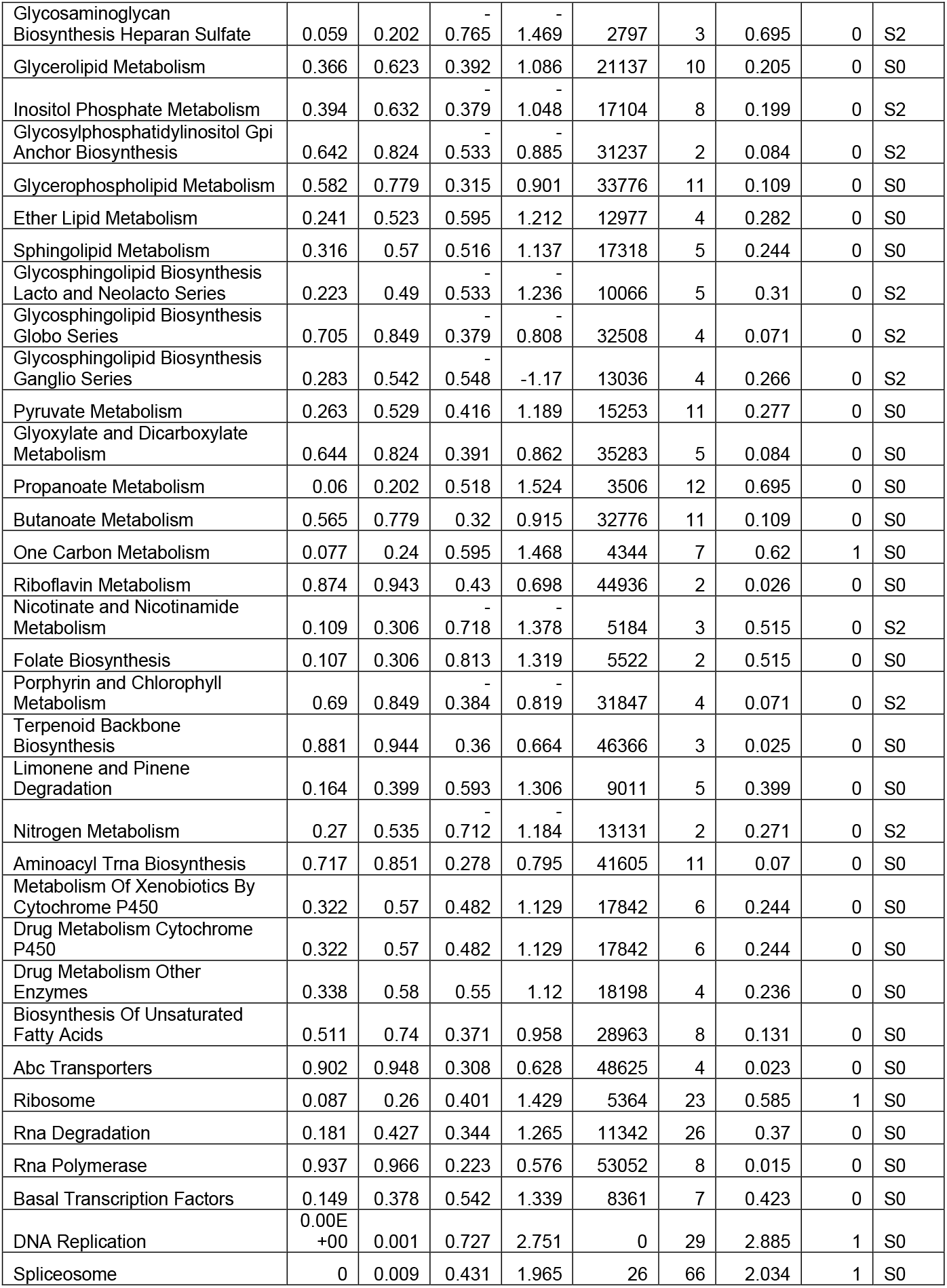
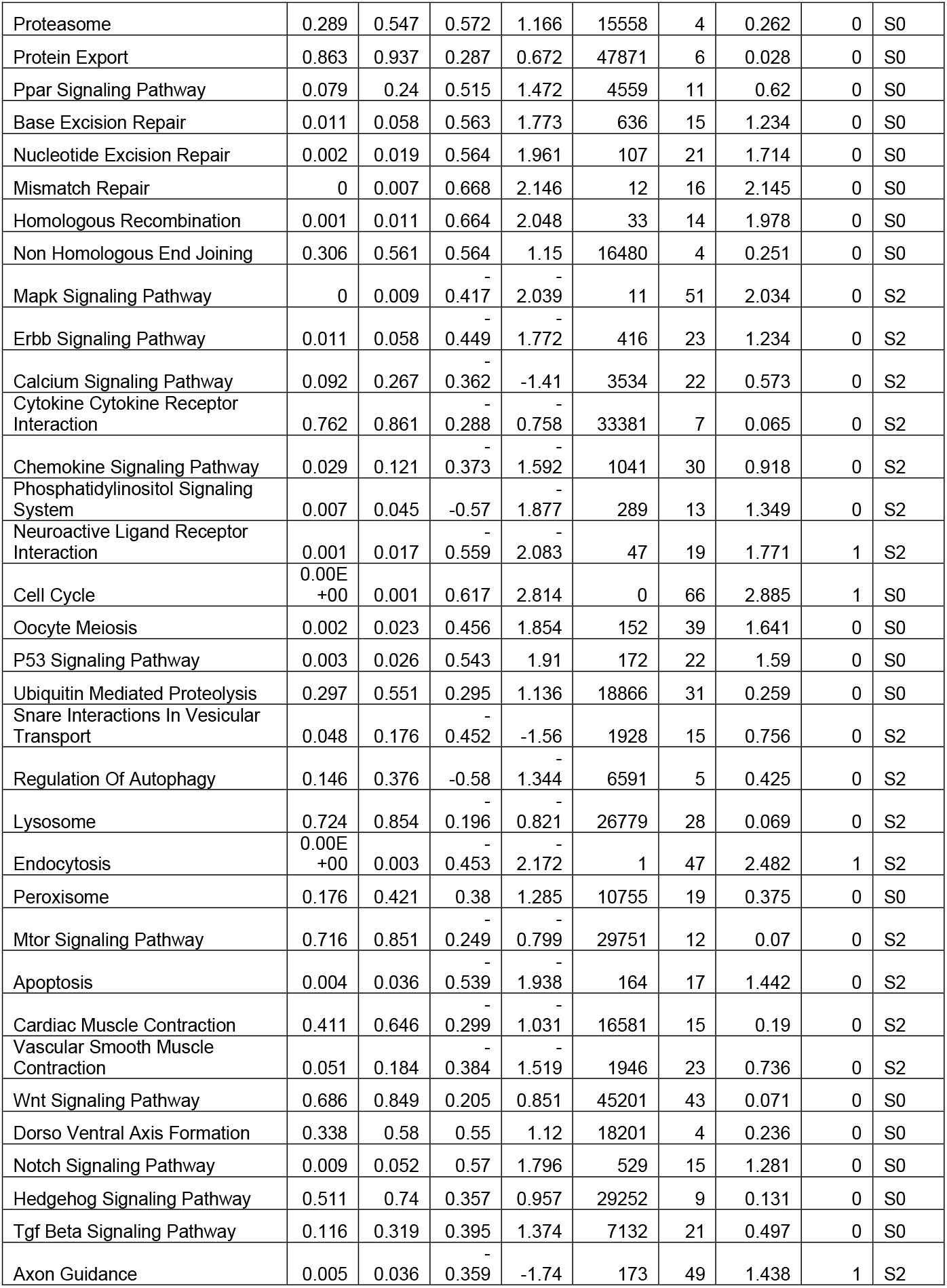
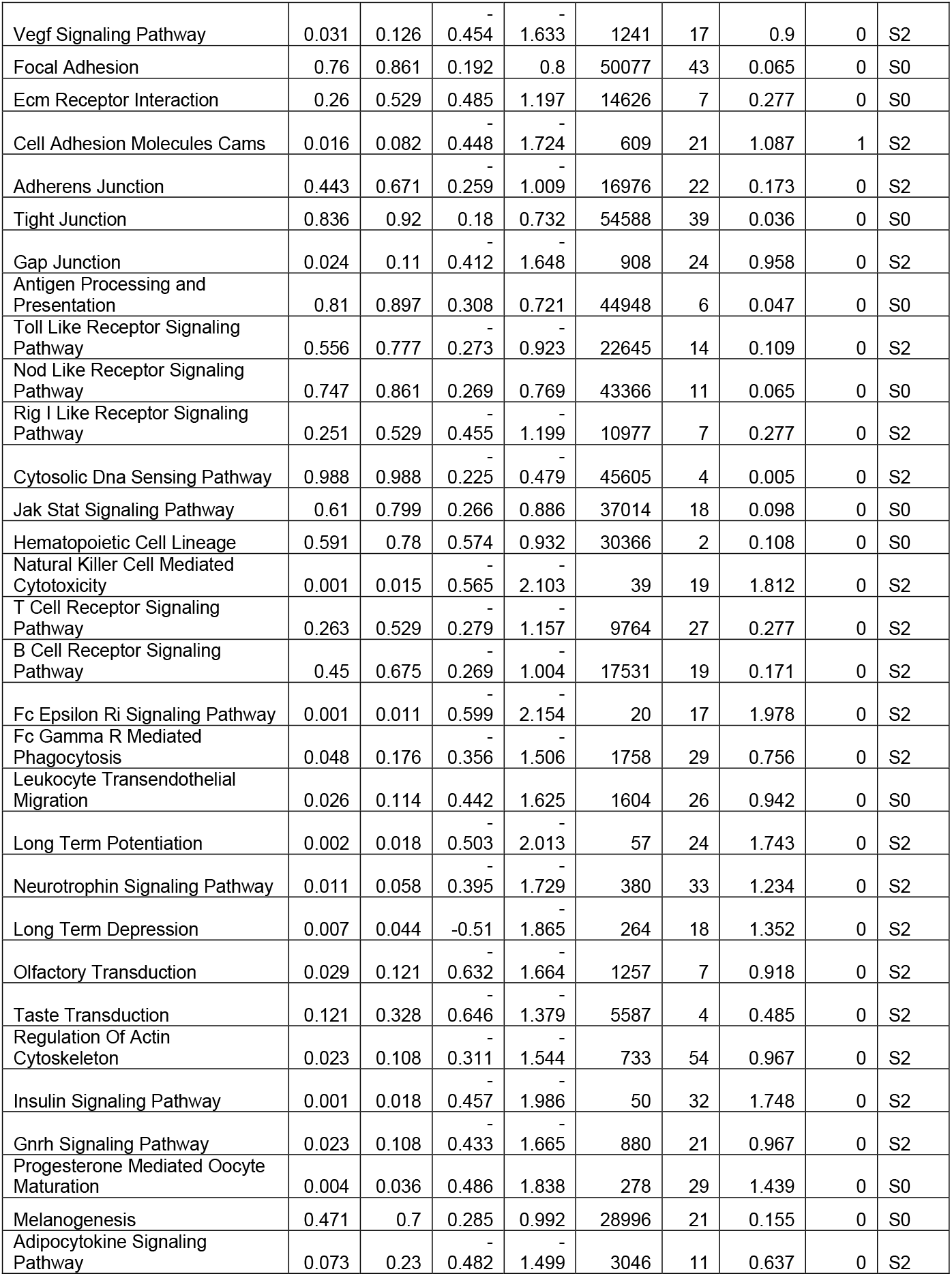
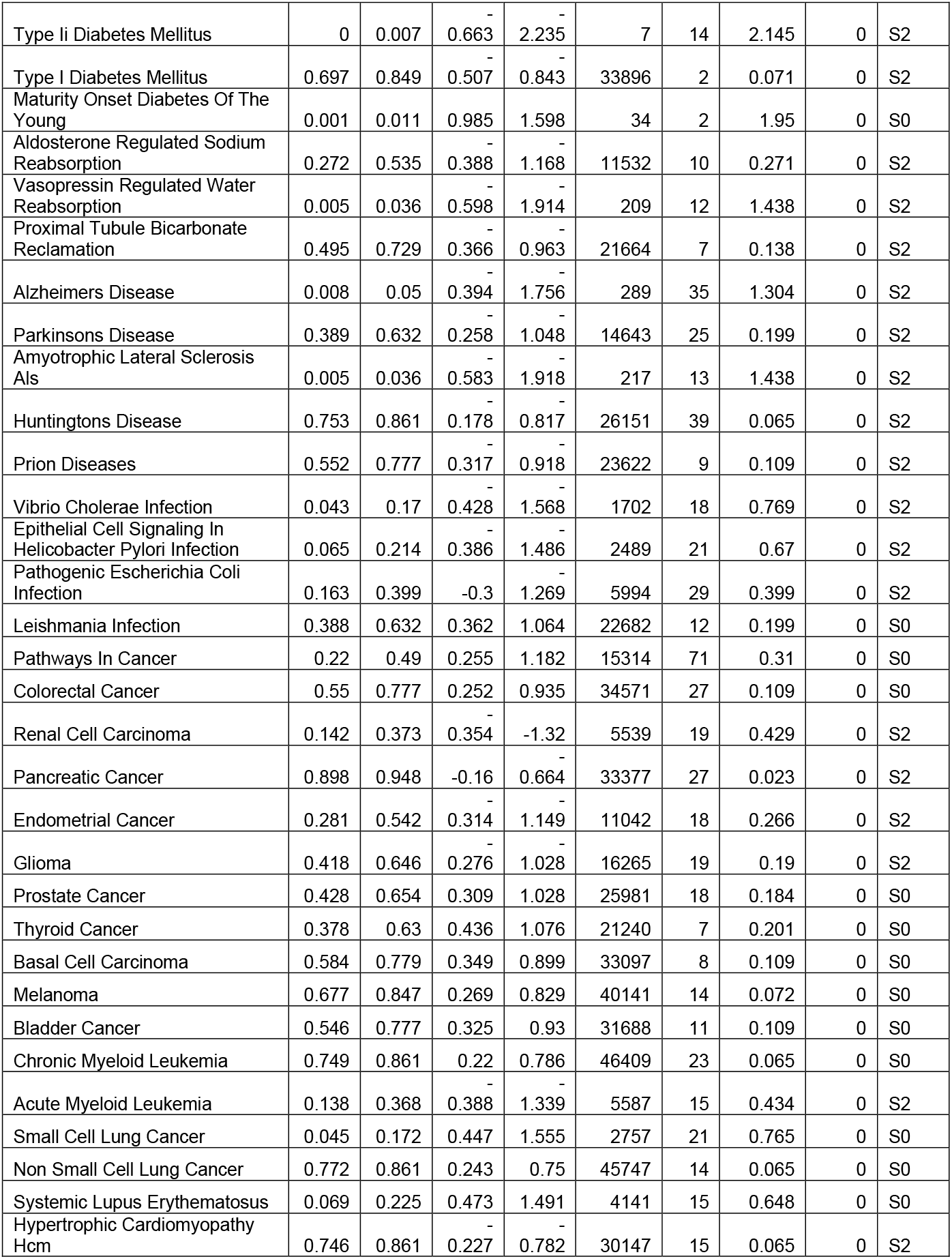
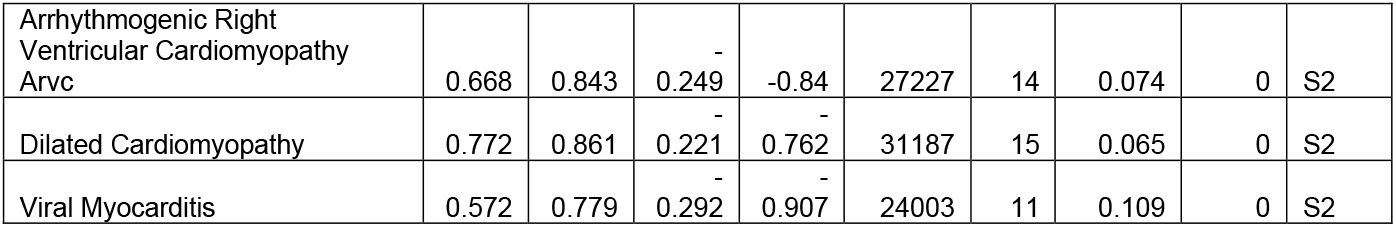
E15.5 S_0_ vs S_2_ fgsea.

**Table S4:** Potential genetic targets for altering cell fate in early neural development.

| Module | Gene Symbol and reference | Description | Known in RPs or neural stem cell context | Summary |
| --- | --- | --- | --- | --- |
| Acetyl-CoA module | Cpt1a (Carnitine Palmitoyltransferase 1A) | Catalyzes the transfer of the acyl group of long-chain fatty acid-CoA conjugates onto carnitine, an essential step for the mitochondrial uptake of long-chain fatty acids and their subsequent beta-oxidation in the mitochondrion. Plays an important role in triglyceride metabolism. | Yes | Required for stem cell maintenance and proper neurogenesis (Knobloch et al., 2017) |
|  | Acadl (Acyl-CoA Dehydrogenase Long Chain) | The protein encoded by this gene belongs to the acyl-CoA dehydrogenase family, which is a family of mitochondrial flavoenzymes involved in fatty acid and branched chain amino-acid metabolism. This protein is one of the four enzymes that catalyze the initial step of mitochondrial beta-oxidation of straight-chain fatty acid. Defects in this gene are the cause of long-chain acyl-CoA dehydrogenase (LCAD) deficiency, leading to nonketotic hypoglycemia. | No | Expressed in SVZ and dentate gyrus (O'Leary et al., 2016) |
|  | Hadh (Hydroxyacyl-CoA Dehydrogenase) | This gene is a member of the 3-hydroxyacyl-CoA dehydrogenase gene family. The encoded protein functions in the mitochondrial matrix to catalyze the oxidation of straight-chain 3-hydroxyacyl-CoAs as part of the beta-oxidation pathway. Its enzymatic activity is highest with medium-chain-length fatty acids. Mutations in this gene cause one form of familial hyperinsulinemic hypoglycemia. The human genome contains a related pseudogene of this gene on chromosome 15. | No | Affects brain development, clinically linked to mental retardation (Yang, He, & Schulz, 2005) |
|  | Acaa2 (Acetyl-CoA Acyltransferase 2) | The encoded protein catalyzes the last step of the mitochondrial fatty acid beta-oxidation spiral. Unlike most mitochondrial matrix proteins, it contains a non-cleavable amino-terminal targeting signal. | No | Its deficiency leads to mental retardation (Fukao, Yamaguchi, Orii, & Hashimoto, 1995) |
|  | Acadm (Acyl-CoA Dehydrogenase Medium Chain) | This gene encodes the medium-chain specific (C4 to C12 straight chain) acyl-Coenzyme A dehydrogenase. The homotetramer enzyme catalyzes the initial step of the mitochondrial fatty acid beta-oxidation pathway. Defects in this gene cause medium-chain acyl-CoA dehydrogenase deficiency, a disease characterized by hepatic dysfunction, fasting hypoglycemia, and encephalopathy, which can result in infantile death. | No | Its deficiency causes encephalopathy (O'Leary et al., 2016) |
|  | Hat1 (Histone Acetyltransferase 1) | The protein encoded by this gene is a type B histone acetyltransferase (HAT) that is involved in the rapid acetylation of newly synthesized cytoplasmic histones, which are in turn imported into the nucleus for de novo deposition onto nascent DNA chains. | No | Maintains acetylation levels (O'Leary et al., 2016) |
|  | Acss1 (Acyl-CoA Synthetase Short Chain Family Member 1) | This gene encodes a mitochondrial acetyl-CoA synthetase enzyme. A similar protein in mice plays an important role in the tricarboxylic acid cycle by catalyzing the conversion of acetate to acetyl CoA. | No | Helps in maintaining acetyl CoA levels (O'Leary et al., 2016) |
| Methylation module | Dnmt3a (DNA Methyltransferase 3 Alpha) | CpG methylation is an epigenetic modification that is important for embryonic development, imprinting, and X-chromosome inactivation. Studies in mice have demonstrated that DNA methylation is required for mammalian development. This gene encodes a DNA methyltransferase that is thought to function in de novo methylation, rather than maintenance methylation. The protein localizes to the cytoplasm and nucleus and its expression is developmentally regulated. | Yes | Deficiency associated with mental retardation (Z. Wu et al., 2012) |
| Glutathione | Gpx8 (Glutathione Peroxidase 8) | GPX8 (Glutathione Peroxidase 8 (Putative)) is a Protein Coding gene. Diseases associated with GPX8 include Anemia, Nonspherocytic Hemolytic, Due To G6pd Deficiency. Among its related pathways are Cellular Senescence (REACTOME) and Glutathione metabolism. Gene Ontology (GO) annotations related to this gene include oxidoreductase activity and peroxidase activity. An important paralog of this gene is GPX7. | Yes | Upregulated when NSCs are exposed to reactive oxidation species (Madhavan, Ourednik, & Ourednik, 2006) |
| Folate module | Dhfr (Dihydrofolate Reductase) | Dihydrofolate reductase converts dihydrofolate into tetrahydrofolate, a methyl group shuttle required for the de novo synthesis of purines, thymidylate, and certain amino acids. While the functional dihydrofolate reductase gene has been mapped to chromosome 5, multiple intronless processed pseudogenes or dihydrofolate reductase-like genes have been identified on separate chromosomes. | Yes | Inhibition leads to neural stem cell differentiation (Fawal et al., 2018) |
|  | Mthfd1 (Methylenetetrahydrofolate Dehydrogenase, Cyclohydrolase And Formyltetrahydrofolate Synthetase 1) | This gene encodes a protein that possesses three distinct enzymatic activities, 5,10-methylenetetrahydrofolate dehydrogenase, 5,10-methenyltetrahydrofolate cyclohydrolase and 10-formyltetrahydrofolate synthetase. Each of these activities catalyzes one of three sequential reactions in the interconversion of 1-carbon derivatives of tetrahydrofolate, which are substrates for methionine, thymidylate, and de novo purine syntheses. | No | Deficiency lead to neural tube defects (J. Wu et al., 2015) |
|  | Tyms (Thymidylate Synthase) | Thymidylate synthase catalyzes the methylation of deoxyuridylate to deoxythymidylate using, 10-methylenetetrahydrofolate (methylene-THF) as a cofactor. This function maintains the dTMP (thymidine-5-prime monophosphate) pool critical for DNA replication and repair. The enzyme has been of interest as a target for cancer chemotherapeutic agents. | No | Deficiency lead to neural tube defects (Wang et al., 2018) |
|  | Shmt1 (Serine Hydroxymethyltransferase 1) | This gene encodes the cytosolic form of serine hydroxymethyltransferase, a pyridoxal phosphate-containing enzyme that catalyzes the reversible conversion of serine and tetrahydrofolate to glycine and 5,10-methylene tetrahydrofolate. This reaction provides one-carbon units for synthesis of methionine, thymidylate, and purines in the cytoplasm. This gene is located within the Smith-Magenis syndrome region on chromosome 17. | No | Deficiency lead to neural tube defects (Beaudin et al., 2011) |
| a-KG module | Phgdh (Phosphoglycerate Dehydrogenase) | This gene encodes the enzyme which is involved in the early steps of L-serine synthesis in animal cells. L-serine is required for D-serine and other amino acid synthesis. The enzyme requires NAD/NADH as a cofactor and forms homotetramers for activity. Mutations in this gene have been found in a family with congenital microcephaly, psychomotor retardation and other symptoms. | Yes | Disruption leads to severe neurodevelopmental defects (Yoshida et al., 2003) |
|  | Psat1 (Phosphoserine Aminotransferase 1) | This gene encodes a member of the class-V pyridoxal-phosphate-dependent aminotransferase family. The encoded protein is a phosphoserine aminotransferase and decreased expression may be associated with schizophrenia. Mutations in this gene are also associated with phosphoserine aminotransferase deficiency. | Yes | Deletion leads to increased stem cell differentiation (Hwang et al., 2016) |
| Cholesterol biosynthesis related | Fdft1 (Farnesyl-Diphosphate Farnesyltransferase 1) | This gene encodes a membrane-associated enzyme located at a branch point in the mevalonate pathway. The encoded protein is the first specific enzyme in cholesterol biosynthesis, catalyzing the dimerization of two molecules of farnesyl diphosphate in a two-step reaction to form squalene. | Yes | Deletion leads to reduced brain size (Coman et al., 2018) |
|  | Hmgcr (3-Hydroxy-3-Methylglutaryl-CoA Reductase) | HMGCR (3-Hydroxy-3-Methylglutaryl-CoA Reductase) is a Protein Coding gene. Diseases associated with HMGCR include Cerebrotendinous Xanthomatosis and Familial Hyperlipidemia. Among its related pathways are Regulation of lipid metabolism by Peroxisome proliferator-activated receptor alpha (PPARalpha) and Statin Pathway. | Yes | Inhibition leads to neurogenesis (Robin et al., 2014) |
|  | Dhcr7 (7-Dehydrocholesterol Reductase) | DHCR7 (7-Dehydrocholesterol Reductase) is a Protein Coding gene. Diseases associated with DHCR7 include Smith-Lemli-Opitz Syndrome and Holoprosencephaly. Among its related pathways are cholesterol biosynthesis I and Vitamin D Metabolism. Gene Ontology (GO) annotations related to this gene include oxidoreductase activity, acting on the CH-CH group of donors, NAD or NADP as acceptor and 7-dehydrocholesterol reductase activity. | Yes | Causes Smith-Lemli-Opitz syndrome (Stelzer et al., 2016) |
| Cancer | Ptprz1 (Protein Tyrosine Phosphatase Receptor Type Z1) | This gene encodes a member of the receptor protein tyrosine phosphatase family. Expression of this gene is restricted to the central nervous system (CNS), and it may be involved in the regulation of specific developmental processes in the CNS. | Yes | Highly upregulated in glioblastoma (Fujikawa et al., 2017) |

